# RNA demethylase FTO uses conserved aromatic residues to recognize the mRNA 5′ cap and promote efficient m^6^A_m_ demethylation

**DOI:** 10.1101/2025.05.09.653100

**Authors:** Brittany Shimanski, Juan F. Marin, Marcin Warminski, Kasun Dilshan Abeyrathne Eluwawalage, Lucas O. Calzini, Riley M. McKeon, Joanna Kowalska, Jacek Jemielity, Jodi A. Hadden-Perilla, Jeffrey S. Mugridge

## Abstract

The RNA demethylase FTO acts as a methyl ‘eraser’ to remove either internal *N*^6^-methyladenosine (m^6^A) or 5′ end *N*^6^-2′-O-dimethyladenosine (m^6^A_m_) modifications on mRNA. FTO has an intrinsic preference and significantly faster demethylation rates *in vitro* for m^6^A_m_ modifications located at the 5′ mRNA cap structure, but the structural basis for FTO’s ability to discriminate m^6^A versus m^6^A_m_ modifications has remained unknown. Here we utilize molecular dynamics simulations of FTO-RNA cap complexes to identify conserved aromatic residues on the surface of FTO involved in 5’ cap recognition. Subsequent mutagenesis and enzymology experiments validate the specificity of these residues in engaging the 5′ cap structure to promote m^6^A_m_ demethylation. We also identify a nonpolar surface on FTO that interacts with the 2′-O-methyl group of m^6^A_m_ to impact demethylation kinetics. This work provides the first structure-level insights into how FTO selectively catalyzes m^6^A_m_ versus m^6^A demethylation on mRNA and advances our understanding of how FTO activity is regulated by diverse mechanisms to help control the epitranscriptome.

## Introduction

*N*^6^-methyladenosine (m^6^A) is the most abundant internal chemical modification of eukaryotic mRNA.^1,2^ Installed co-transcriptionally by the METTL3/14 methyltransferase ‘writer’ complex, m^6^A can be found throughout the length of mRNA transcripts and is enriched near stop codons and in the 3′ UTR.^3,4^ This widely important modification has impacts on mRNA splicing,^5–7^ export,^8,9^ translation,^10–13^ and stability.^14–16^ Notably, m^6^A plays a crucial role in the regulation of mammalian stem cell differentiation^17^ and is involved in controlling the expression of human oncogenes and tumor suppressor genes.^18–20^ Another highly abundant mRNA methyl modification, *N*^6^,2′-O-dimethyladenosine (m^6^A_m_), is found exclusively at the 5′ end of eukaryotic mRNAs in up to 30% of mammalian 5′ mRNA caps.^21^ m^6^A_m_ is installed by the writer enzyme CAPAM (or PCIF1), which carries out *N*^6^-methylation of 2′-O-methyladenosine (A_m_) modifications.^22–25^ While the functions of m^6^A_m_ modifications remain incompletely defined and may vary cellular context, recent work suggests these 5′ cap modifications may influence mRNA stability, translation and transcription, whereas m^6^A_m_ modifications in snRNA may help regulate snRNA biogenesis and splicing.^26–33^ Together, the highly abundant methyl modifications m^6^A and m^6^A_m_ help control mRNA fate and function across the transcriptome.

Fat Mass and Obesity Associated Protein (FTO) is an Fe(II)/2-oxoglutarate-dependent dioxygenase enzyme that acts as an m^6^A demethylase or ‘eraser’, converting m^6^A or m^6^A_m_ modifications to A or A_m_, respectively, within cellular mRNA transcripts.^34–37^ Along with other AlkB homologs in this family (e.g. ALKBH5), FTO allows for reversible m^6^A and m^6^A_m_ modification and the dynamic regulation of RNA methylation. Although FTO likely only acts on a subset of m^6^A/m^6^A_m_ modifications found on RNA,^33,38,39^ its m^6^A/m^6^A_m_ demethylation activity is linked to splicing,^7,27,40,41^ cellular metabolism,^42–46^ and mammalian development.^47–49^ Furthermore, FTO and its methyl eraser functions have close links to the progression of human cancers, such as acute myeloid leukemia,^50,51^ colorectal cancer,^52,53^ and glioblastoma.^54,55^ More broadly, FTO can promote oncogenesis and immune evasion in different cancer contexts.^56–58^

Although FTO has the ability to demethylate both internal m^6^A and 5′ cap m^6^A_m_ RNA modifications, previous studies have shown that FTO has a strong intrinsic preference *in vitro* for m^6^A_m_ demethylation^26^ and more recent studies call into question whether FTO truly targets m^6^A modifications in the cell.^59,60^ Remarkably, FTO has ∼100-fold greater catalytic efficiency for demethylating m^6^A_m_ compared to m^6^A modifications on RNA oligonucleotides.^26^ Furthermore, in model RNA substrates where the 5′ *N*^7^-methylguanosine (m^7^G) and/or triphosphate groups of the eukaryotic cap are removed, FTO-mediated demethylation of m^6^A_m_ is significantly impaired, suggesting that FTO somehow specifically recognizes the 5′ cap structure to promote m^6^A_m_ demethylation. An existing structure of 6mA-ssDNA in complex with FTO provides information about how the m^6^A base is accommodated in the FTO active site and how DNA (and likely RNA) binds a basic cleft on the surface of FTO.^37^ However, to date there is no structural or biophysical information about how FTO preferentially demethylates 5′ m^6^A_m_ modifications, how FTO might specifically recognize the eukaryotic 5′ cap structure and m^7^G or m^6^A_m_ bases, or the general mechanism(s) by which FTO selectively catalyzes removal of m^6^A_m_ versus m^6^A modifications on RNA.

Here we combine molecular dynamics (MD) simulations of FTO-m^6^A_m_ cap complexes with mutagenesis and enzymology experiments to identify conserved aromatic residues on the surface of FTO involved in 5′ mRNA cap recognition and selective m^6^A_m_ demethylation. We also identify nonpolar residues located just outside of the FTO active site that interact with the m^6^A_m_ 2′-O-methyl group to further mediate m^6^A_m_ demethylation. These experiments provide the first atomic-level information about how 5′ cap recognition promotes m^6^A_m_ demethylation by FTO and give new insights into the mechanisms controlling FTO-mediated mRNA demethylation selectivity.

## Results

### Molecular dynamics simulations identify conserved aromatic FTO surface residues involved in FTO-m^7^G cap interactions

Previous biochemical studies have shown that FTO more efficiently demethylates 5′ m^6^A_m_ modifications as compared to internal m^6^A modifications on RNA.^26^ Additionally, removing the m^7^G portion of the 5′ mRNA cap structure on model RNA substrates results in a 50% reduction in m^6^A_m_ demethylation activity, and further removal of the cap triphosphate group reduces demethylation by another 40%. These data suggest that there must be FTO residues that selectively recognize the m^7^G and triphosphate groups adjacent to m^6^A_m_ and that these surfaces help to recruit and/or position 5′ m^6^A_m_ modifications for demethylation.^26^ To elucidate which residues and surfaces of FTO are interacting with the 5′ mRNA cap structure, we performed microsecond MD simulations of FTO in complex with the small 5′ cap molecule m^7^Gppp(m^6^A_m_)pG (**Figure 1A**) in triplicate. Molecular mechanics/generalized Born surface area (MM/GBSA) calculations were applied to the MD trajectories to identify residues contributing to m^7^Gppp(m^6^A_m_)pG binding affinity **(Supplementary Figure 1)**. Of these, three conserved aromatic residues on the surface of FTO were observed to make high-frequency contacts (≤ 3.5 Å) with the 5′ m^7^G base or ribose (**Figure 1B, Supplementary Figure 2**): Y220 (**Figure 1C**), H232 (**Figure 1D**), and W278 (**Figure 1E**).

**Figure 1.**
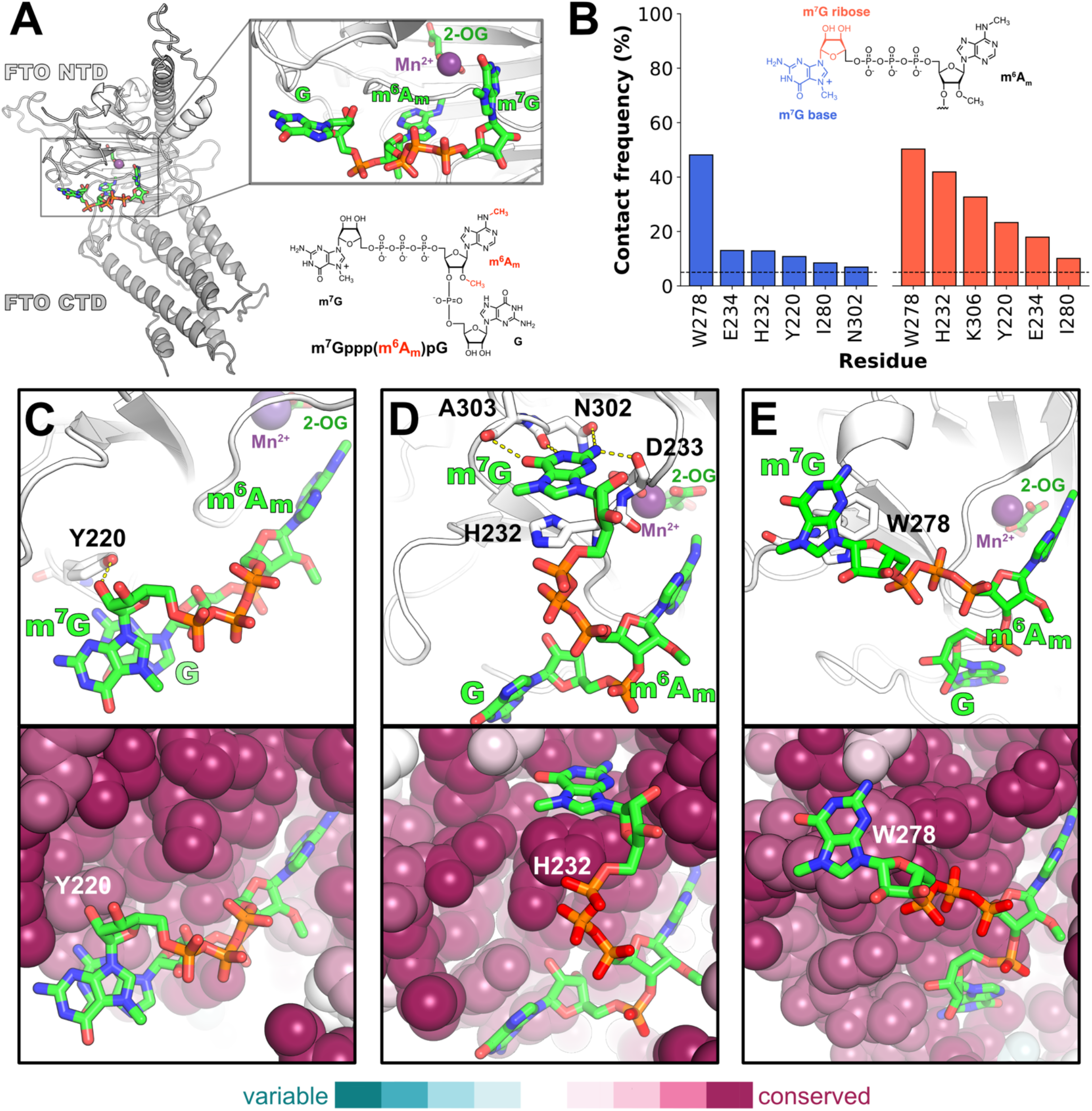
MD reveals FTO residues with high-frequency FTO-m^7^G interactions. **(A)** To provide a starting point for MD simulations, an m^7^Gppp(m^6^A_m_)pG cap structure was positioned on FTO by aligning the m^6^A_m_ base into the FTO active site based on previously determined 6mA-ssDNA-FTO structure PDB 5ZMD. **(B)** m^7^G-FTO residue contacts ≤ 3.5 Å with > 5% interaction frequency from MD simulations on the microsecond timescale. Only FTO residues with favorable contributions to binding affinity (ΔG ≤ −0.25 kcal/mol) are considered. Per-residue MM/GBSA decomposition for each simulation replicate is provided in **Supplementary Figure 1. (C)** Representative MD frame showing hydrogen-bonding interaction (dashed yellow line) between FTO residue Y220 and m^7^G ribose of the cap structure (top). ConSurf^61,62^ analysis shows this is a strongly conserved surface of FTO (bottom). **(D)** Representative MD frame showing m^7^G stacking with conserved FTO residue H232 and hydrogen bonding with N302, A303, and D233 side chain and backbone atoms on FTO surface; hydrogen bonds are shown as dashed yellow lines (top). In this binding mode m^7^G interacts with a highly conserved surface of FTO (bottom). **(E)** Representative MD frame showing m^7^G stacking on strongly conserved FTO residue W278.

**Figure 2.**
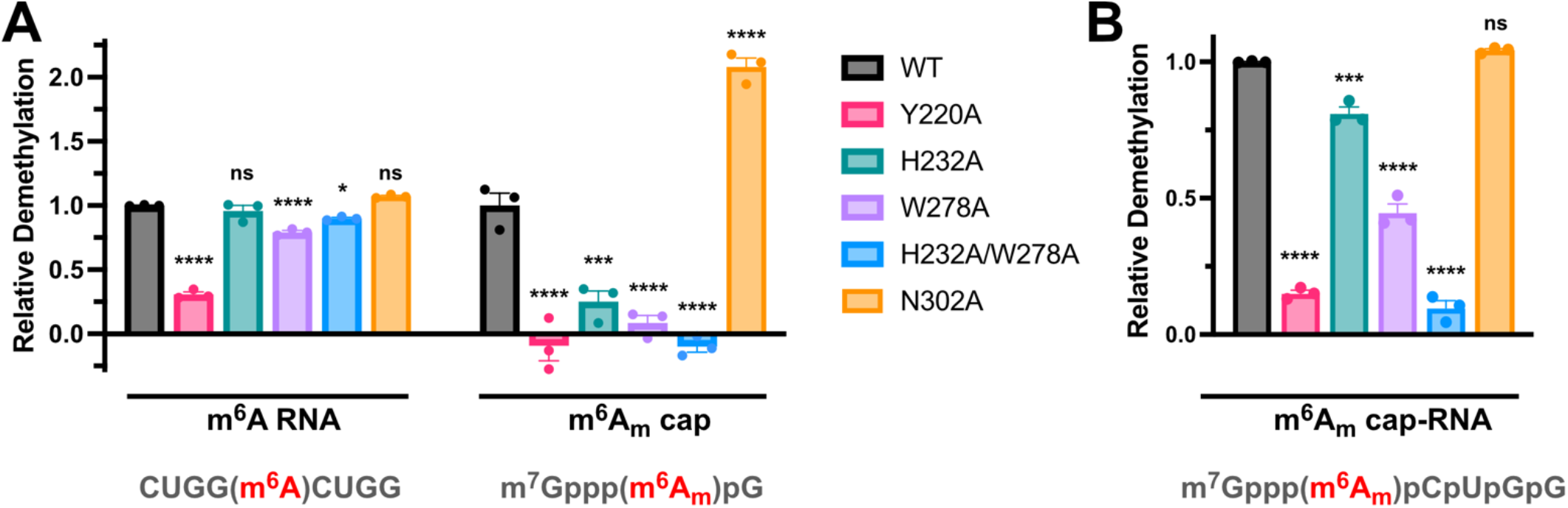
Enzymatic assays validate FTO residues important for m^6^A_m_ demethylation. **(A)** Mutation of conserved aromatic residues in FTO identified in MD simulations as m^7^G interaction sites show altered m^6^A_m_ demethylation activity. Endpoint demethylation activity assays were conducted in triplicate with 2 µM CUGG(m^6^A)CUGG (m^6^A-RNA) or m^7^Gppp(m^6^A_m_)pG (m^6^A_m_ cap) and 1 µM FTO enzyme. At 0 and 150 minutes, reaction samples were quenched with EDTA, digested to single nucleosides with RNAase cocktail, and the relative amounts of m^6^A or m^6^A_m_ were quantified by UHPLC-MS. **(B)** Mutation of conserved aromatic residues in FTO identified as m^7^G interaction sites show reduced m^6^A_m_ demethylation activity with a longer m^6^A_m_ cap RNA substrate. Endpoint demethylation activity assays were conducted with 2 µM m^7^Gppp(m^6^A_m_)pCpUpGpG (m^6^A_m_ cap-RNA) and 0.1 µM FTO enzyme in triplicate. At 0 and 150 minutes, reaction samples were quenched with EDTA, decapped, digested to single nucleosides with RNAase cocktail, and the relative amounts of m^6^A_m_ were quantified by UHPLC-MS. For both (A) and (B), statistical significance was calculated using a one-way ANOVA with Dunnett’s multiple comparisons test (ns p > 0.05, * p ≤ 0.05, ** p ≤ 0.01, *** p ≤ 0.001, **** p ≤ 0.0001); errors shown as SEM.

MD simulations show that Y220 can interact with the m^7^G ribose, forming hydrogen bonds through its hydroxyl group (**Figure 1C**). The phenol ring of Y220 was not observed to participate in guanine base stacking unless m^6^A_m_ adopted a shallow, catalytically ineffective position in the FTO active site (**Supplementary Figure 3**). The H232 imidazole ring can form a cation-π / stacking interaction with the positively charged m^7^G, while the base makes simultaneous hydrogen bond contacts with backbone atoms of FTO residues N302, A303, and D233, as well as the N302 side chain (**Figure 1D**). Alternatively, H232 can make CH-π or hydrogen bond contacts with the m^7^G ribose or interact with the triphosphate bridge (ppp). H232 is located on the surface of FTO, immediately adjacent to metal-binding residues in the FTO active site and was treated as neutral during MD simulations. Finally, W278, located on a flexible loop of FTO, can form a cation-π / stacking interaction with the positively charged m^7^G base (**Figure 1E**).

**Figure 3.**
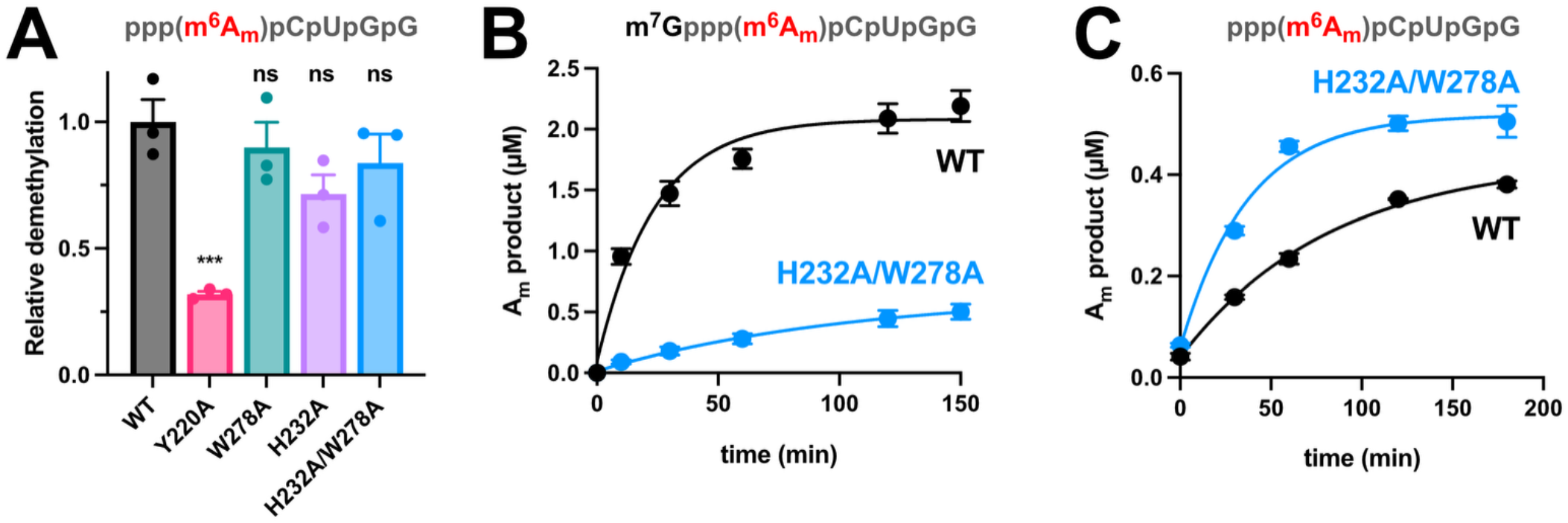
Demethylation assays with truncated caps implicate FTO residues H232 and W278 in selective m^7^G recognition. **(A)** Endpoint demethylation assays comparing the relative m^6^A_m_ demethylation activity for WT versus mutant FTO constructs with the m^7^G-lacking ppp(m^6^A_m_)pCpUpGpG substrate. Triplicate demethylation assays were carried out as in **Figure 2**, with 2 µM ppp(m^6^A_m_)pCpUpGpG substrate and 1 µM FTO enzyme. FTO Y220A has impaired m^6^A_m_ demethylation activity on the truncated, m^7^G-less substrate, whereas H232A, W278A, and H232A/W278A mutants have no significant defect and similar activity to WT. Statistical significance was calculated using a one-way ANOVA with Dunnett’s multiple comparisons test (ns p > 0.05, *** p ≤ 0.001); data are shown as mean values ± SEM (n = 3). **(B)** Time courses comparing kinetics of A_m_ product formation for WT versus H232A/W278A FTO with m^6^A_m_ cap substrate m^7^Gppp(m^6^A_m_)pCpUpGpG show that the FTO H232A/W278A double mutant has significantly impaired demethylation kinetics compared to WT. **(C)** Time courses comparing kinetics of A_m_ product formation for WT versus H232A/W278A FTO with truncated m^6^A_m_ cap substrate ppp(m^6^A_m_)pCpUpGpG, which lacks the m^7^G group, show that the FTO H232A/W278A double mutant has no defect in demethylation kinetics compared to WT.

Each of these three aromatic FTO residues (Y220, H232, W278) are strongly conserved (**Figure 1C-E**, bottom) and can engage with the 5′ cap structure in multiple binding modes, including modes that simultaneously involve two of the aromatic residues, for example H232 interacting with the m^7^G ribose while W278 stacks with the m^7^G guanosine base (**Supplementary Figure 4A**), or W278 and Y220 sandwiching m^7^G (**Supplementary Figure 4B**).

**Figure 4.**
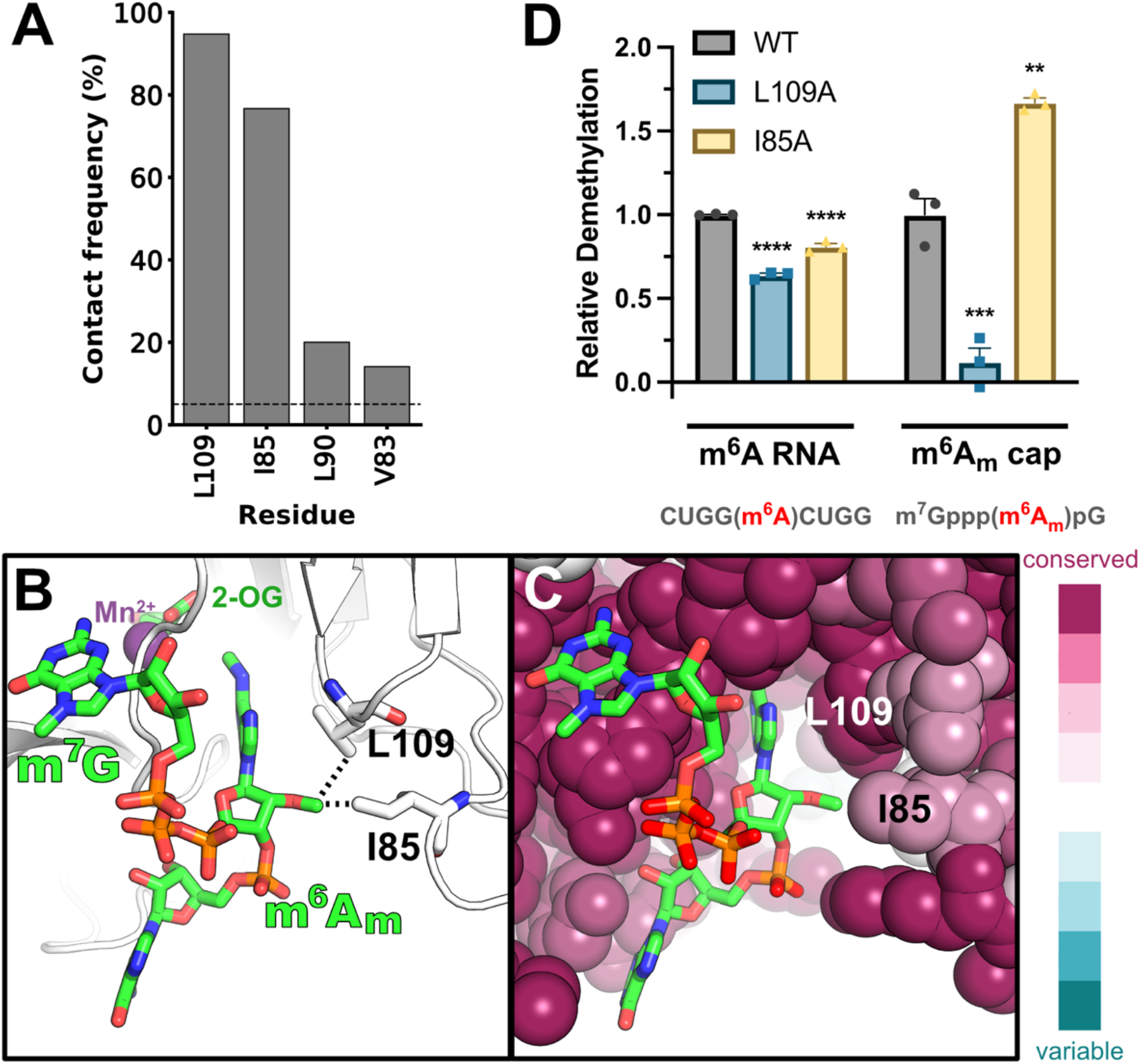
Nonpolar FTO residues help recognize the A_m_ 2′-O-methyl group to promote m^6^A_m_ demethylation. **(A)** Graph showing high-frequency A_m_ 2′-O-methyl-FTO residue contacts ≤ 3.5 Å from MD simulations on the microsecond timescale. Only FTO residues with favorable contributions to binding affinity (ΔG ≤ −0.25 kcal/mol) are considered. Per-residue MM/GBSA decomposition for each simulation replicate is provided in **Supplementary Figure 1. (B)** Representative frame from MD simulations showing close interactions (dashed lines) of the m^6^A_m_ 2′-O-methyl group with the nonpolar FTO residues L109 and I85. **(C)** L109 and I85 form a strongly conserved nonpolar surface located just outside the FTO active site pocket. **(D)** FTO L109A and I85A mutations show altered demethylation activity for m^6^A_m_ cap compared to internal m^6^A-RNA substrates. Endpoint demethylation activity assays were conducted in triplicate with 2 µM CUGG(m^6^A)CUGG or m^7^Gppp(m^6^A_m_)pG and 1 µM FTO enzyme. Statistical significance was calculated using a one-way ANOVA with Dunnett’s multiple comparisons test (** p ≤ 0.01, *** p ≤ 0.001, **** p ≤ 0.0001); errors shown as SEM.

### Mutations to conserved aromatic surface residues on FTO specifically impair m^6^A_m_ demethylation

To experimentally validate the FTO-m^7^G interactions identified in the MD simulations described above, we mutated key, high-frequency m^7^G-interacting FTO residues to alanine and compared WT versus mutant demethylation activities on both an internal m^6^A-containing RNA substrate and the model cap substrate m^7^Gppp(m^6^A_m_)pG (**Figure 2A**). Model cap substrates were used in these initial screening experiments to maximize the likelihood that defective m^7^G or cap recognition in mutant FTO constructs results in a strong and observable effect on demethylation activity. LC-MS-based demethylation activity assays were conducted on the panel of FTO mutants to quantify relative demethylation activity after 150 minutes compared to WT FTO. These biochemical assays showed that FTO Y220A mutation negatively impacted both m^6^A and m^6^A_m_ demethylation; H232A, N302A, W278A, and H232A/W278A mutations had little to no defect on demethylation of 9-mer m^6^A RNA substrate (**Figure 2A**, ‘m^6^A-RNA’); while H232A, W278A, and H232A/W278A mutants all significantly impaired demethylation of m^6^A_m_ in the model cap substrate (**Figure 2A**, ‘m^6^A_m_ cap’). Time courses with FTO mutants showed similar defects in m^6^A_m_ demethylation kinetics with H232A and W278A mutants (**Supplementary Figure 5**). Surprisingly, N302A had ∼2-fold higher activity on m^6^A_m_ cap compared to WT FTO. Notably, MD simulations showed that in the absence of m^7^G interactions, N302 and H232 frequently form hydrogen bonds with each other; loss of the N302 side chain may therefore increase accessibility of H232 for cap recognition, consistent with increased activity in the N302A mutant.

We next tested the relative m^6^A_m_ demethylation activities for WT versus mutant FTO constructs on the longer capped model RNA substrate m^7^Gppp(m^6^A_m_)pCpUpGpG, which more closely mimics a capped RNA. With this longer m^6^A_m_-containing substrate we observed a significant defect in demethylation activity with Y220A mutation, a small but measurable defect on m^6^A_m_ demethylation for mutation of FTO H232, and, very similar to the results above with the shorter cap substrate, large defects in activity with W278A and H232A/W278A mutants (**Figure 2B**). Enzymological analysis suggests that the defect in m^6^A_m_ demethylation observed with double mutant H232A/W278A is predominantly a *k*_cat_, rather than *K*_M_, effect (**Supplementary Figure 6**). Together, these kinetic data show that mutation of FTO residues H232 and W278 significantly impair FTO-mediated demethylation activity of capped m^6^A_m_-containing RNAs, but not internal m^6^A-containing RNAs, suggesting these two conserved FTO surface residues play a key role in recognizing the 5′ cap to promote efficient m^6^A_m_ demethylation catalysis. While Y220A mutation also impairs m^6^A_m_ demethylation, it has similar effects on m^6^A demethylation, suggesting that this mutation more generally impairs FTO activity and/or RNA binding.

### FTO residues H232 and W278 specifically recognize m^7^G of cap

The MD simulations described above identified key conserved residues on the surface of FTO that have high-frequency interactions with the m^7^G group of the RNA 5′ cap, and our biochemical validation experiments showed that mutation of these residues impairs m^6^A_m_ demethylation. We next asked if removal of the m^7^G group from our model cap-RNA substrates eliminates the relative defect in m^6^A_m_ demethylation activity between FTO WT and mutants – as would be expected for FTO residues that specifically and exclusively recognize m^7^G. In demethylation assays with ppp(m^6^A_m_)pCpUpGpG (ppp-m^6^A_m_-RNA, **Figure 3A**), we found that while FTO Y220A still has a strong defect in demethylation activity, H232A, W278A, and H232A/W278A mutants no longer have significant defects in activity compared to WT on this substrate that lacks m^7^G. To further confirm this, we compared time courses for m^6^A_m_ demethylation for WT versus H232A/W278A FTO with the normal m^7^G-containing m^6^A_m_-RNA substrate (**Figure 3B**) and with the m^7^G-lacking ppp-m^6^A_m_-RNA substrate (**Figure 3C**). Consistent with the endpoint demethylation assays, these experiments show that the FTO H232A/W278A mutant has a dramatic defect in m^6^A_m_ demethylation activity *only when m*^*7*^*G is present* in the substrate; removal of m^7^G lowers the overall demethylation activity, but results in WT and H232A/W278A having similar m^6^A_m_ demethylation rates. Together, these data strongly suggest that FTO H232 and W278 are responsible for selective recognition of the m^7^G group of the 5′ cap structure to promote m^6^A_m_ demethylation, because mutation of these residues results in demethylation defects only on m^7^G-containing m^6^A_m_ cap substrates – but not on internal m^6^A-containing substrates or m^6^A_m_ cap substrates that lack m^7^G.

### MD simulations and kinetic assays identify FTO residues important for recognition of the 2′-O-methyl group in m^6^A_m_

Similar to m^7^G, the 2′-O-methyl group on adenosine (A_m_) in the m^6^A_m_ cap structure has also been shown to be important for promoting FTO-mediated demethylation of m^6^A_m_.^26^ *In vitro* activity studies indicate that absence of the A_m_ 2′-O-methyl group in model m^7^Gppp(m^6^A)-RNA substrates results in a 50% decrease in FTO’s ability to demethylate the 5′ m^6^A group as compared to m^6^A_m_. However, there is no structural information about how FTO recognizes the A_m_ 2′-O-methyl group to help promote m^6^A_m_ demethylation. To explore this FTO-m^6^A_m_ interaction, we surveyed A_m_ methyl-FTO contacts ≤ 3.5 Å over all frames in the FTO-m^6^A_m_ MD simulations and identified two FTO residues, L109 and I85, that contact the A_m_ 2′-O-methyl group with very high frequencies (**Figure 4A, Supplementary Figure 7**). These strongly conserved nonpolar residues are located just outside the active site pocket and provide a hydrophobic surface on which the A_m_ 2′-O-methyl group frequently rests throughout the majority of the MD simulations (**Figure 4B,C**). Demethylation activity assays comparing WT FTO to L109A or I85A mutants revealed that while both of these mutations resulted in only small defects to m^6^A demethylation on an internal m^6^A-RNA, L109A resulted in a nearly 90% reduction in m^6^A_m_ cap demethylation activity (**Figure 4D**). Surprisingly, the FTO I85A mutation resulted in a ∼50% increase in m^6^A_m_ cap demethylation relative to WT, suggesting this mutation may open up additional space on this surface or shallow pocket to accommodate the A_m_ 2′-O-methyl group. Together, these experiments suggest that this nonpolar surface on FTO (L109, I85) plays a key role in interacting with the 2′-O-methyl group of the m^6^A_m_ base to help mediate N^6^ demethylation.

## Discussion

Previous biochemical studies have shown that FTO has a clear intrinsic preference for ‘erasing’ m^6^A_m_ modifications located at the 5′ end of mRNA, as compared to internal m^6^A modifications.^26^ While the 6mA-ssDNA-bound FTO structure determined in 2019 provides the structural basis for m^6^A base recognition in the active site,^37^ it remains entirely unknown how FTO recognizes the eukaryotic 5′ cap structure and m^6^A_m_ 2′-O-methyl group to promote efficient 5′ m^6^A_m_ demethylation on mRNA. In this work, using MD simulations we identified conserved, aromatic FTO residues H232 and W278 as key surfaces for high-frequency interaction with the m^7^G base of the m^6^A_m_ cap structure. Likewise, MD simulations show that nonpolar FTO residues L109 and I85 form a key interaction surface for the m^6^A_m_ 2′-O-methyl group. *In vitro* activity assays validate the structural predictions from MD, and show that mutating FTO residues H232, W278, or L109 to alanine selectively impairs 5′ m^6^A_m_ demethylation, with only minimal effects on internal m^6^A demethylation. Although FTO residue Y220 was also identified as a high-frequency site of cap interaction, the majority of MD conformations with Y220-m^7^G interactions show either Y220 hydrogen bonding with the m^7^G ribose, or the m^6^A_m_ base in a catalytically-incompetent position when the m^7^G base stacks with Y220; biochemical experiments show that Y220 mutation impairs both m^6^A and m^6^A_m_ demethylation, suggesting Y220 may help promote interactions with RNA that are not cap-specific. Together, our MD and biochemical data provide the first atomic-level insights into how FTO recognizes the m^7^G base (H232/W278) and the 2′-O-methyl group (L109/I85) of 5′ m^6^A_m_-modified mRNAs.

The MD simulations and enzymology data both support the conclusion that FTO surface residues H232 and W278 contribute to m^7^G binding, cap recognition and selective m^6^A_m_ demethylation. It appears that m^7^G and the 5′ cap dynamically sample multiple surfaces of FTO, as we observe in the MD simulations, and this leads to the importance of both H232 and W278 for cap recognition. Both of these residues can interact simultaneously with the cap, with W278 engaging the m^7^G base while H232 engages the m^7^G ribose (**Supplementary Figure 4A**), or either residue can independently engage the m^7^G base (**Figure 1D,E**), providing a potentially redundant mechanism that preserves cap binding across multiple conformational states. Additionally, W278 is located on a flexible loop of FTO, and MD suggests that W278 can in some instances come close enough to H232 for both of these residues to stack simultaneously with the m^7^G base of the cap (**Supplementary Figure 8A**). Therefore, another possibility consistent with our experimental data is that FTO H232 and W278 could sandwich the m^7^G base, making dual cation-π interactions that specifically recognize the 5′ cap structure. This mode of recognition would be very similar to how canonical cap-binding proteins, such as the cap-binding complex (CBC; CBP20/80),^63^ eIF4E,^64^ or Dcp2,^65,66^ recognize the mRNA cap (**Supplementary Figure 8B**) and would be consistent with specific m^7^G recognition by FTO.

Furthermore, m^7^G binding modes where m^7^G stacks with H232 (as in **Figure 1D**) may explain a puzzling piece of biochemical data from the literature. Recently it was shown that *N*^2,2,7^-trimethylguanosine (m^2,2,7^G)-capped RNAs, like those utilized during splicing, are entirely resistant to m^6^A_m_ demethylation by FTO;^27^ this is surprising because removal of m^7^G entirely only reduces m^6^A_m_ demethylation activity by 50%.^26^ But when m^7^G stacks with H232 as shown in **Figure 1D**, the N2 nitrogen of guanosine points directly toward the key metal binding active site residue D233. Dimethylation of the N2 atom, as in m^2,2,7^G, would sterically clash with D233 and be predicted to disrupt metal binding and inactivate FTO, consistent with the experimental observation that m^2,2,7^G-capped RNAs have no detectable m^6^A_m_ demethylation activity with FTO.

Our combined MD and biochemical studies reveal key, conserved residues on the surface of FTO that recognize the eukaryotic 5′ cap structure to promote efficient m^6^A_m_ demethylation on mRNA. These results provide a structural basis for FTO’s ability to distinguish 5′ m^6^A_m_ modifications from internal m^6^A RNA modifications, and suggest that dynamics of FTO residues and/or the 5′ cap may be important for selective cap recognition. Finally, we propose that engineered FTO mutants that specifically impair m^6^A_m_ demethylation, but minimally affect m^6^A demethylation (e.g., H232A/W278A, H232A/W278A/L109A), could be used as separation-of-function mutants^67^ in cell-based studies to dissect and understand the different biological roles of FTO-mediated m^6^A_m_ versus m^6^A demethylation across the transcriptome.

## Methods

### Model construction

An all-atom model of FTO in complex with the small 5′ cap molecule m^7^Gppp(m^6^A_m_)pG was developed based on PDB 5ZMD.^37^ Missing loop segments in FTO were modeled using AlphaFold2.^68^ The m^7^Gppp(m^6^A_m_)pG cap structure was built with xleap^69^ using residue templates from OL3.^70^ The m^6^A_m_ base was aligned into the FTO catalytic pocket based on the 6mA orientation in the FTO-6mA-ssDNA structure. Crystallographic water molecules were maintained. PDB2PQR^71^ was used to add hydrogen atoms to FTO appropriate for pH 7.0. Local ions were placed around the complex with cionize,^72^ and the complex was immersed in a 125 Å^3^ box of OPC^73^ water containing 150 mM NaCl.

### Model parameterization

FTO was treated with the ff19SB^74^ force field. Manganese(II) coordination was parameterized using MCPB.py^75^ from AmberTools25.^76^ The 2-oxoglutarate cofactor was treated with gaff2^77^ parameters and AM1-BCC charges assigned by antechamber.^69^ A custom residue was constructed for m^7^Gppp(m^6^A_m_)pG. Parameters were taken from OL3^70^ based on analogy to a GAG sequence, applying updated phosphate oxygen radii.^78^ Methyl and triphosphate parameters were taken from gaff2.^77^ Missing parameters for the methyl-purine moiety in m^7^G (two angles and an improper dihedral), as well as residue charges, were taken from the modified nucleoside force field,^79^ which covers methylated RNA.

### Molecular dynamics simulations

MD simulations in the isothermal-isobaric (NPT) ensemble were performed with pmemd.cuda (SPFP precision model) in AMBER24.^80^ The system was brought to a local energy minimum using 2,500 steepest descent cycles, then 2,500 conjugate gradient cycles. The temperature was raised from 60 K to 310 K over a period of 5 ns, then production sampling was carried out for 1 μs, saving frames every 1 ps. Dynamics were propagated with a timestep of 2.0 fs, enabled by constraining bonds to hydrogen with the SHAKE algorithm. Non-bonded interactions were partitioned into long-and short-range components based on a cutoff of 8.0 Å. System temperature was maintained at a target of 310 K with the Langevin thermostat using a collision frequency of 1.0 ps^-1^, reinitializing the random number seed upon each simulation restart to avoid synchronization artifacts.^81^ System pressure was maintained at a target of 1.0 bar via isotropic scaling with the Berendsen barostat using a relaxation time of 1.0 ps.

### Trajectory analysis

MD simulation trajectories were analyzed using VMD 1.9.4.^72^ Pairwise contacts between FTO and m^7^Gppp(m^6^A_m_)pG were defined as being atoms ≤3.5 Å to filter for the most relevant interactions. Only residues contributing meaningfully to FTO-m^7^Gppp(m^6^A_m_)pG binding affinity were considered for contact analysis. Binding affinity was estimated using molecular mechanics/generalized Born surface area (MM/GBSA)^82,83^ calculations, performed using mmpbsa.py^84^ in AmberTools25.^76^ MM/GBSA used the Onufriev et al. GB model II (igb=5)^85^ with the single-trajectory protocol, in which the complex, receptor, and ligand energies are evaluated from the same trajectory. Ionic strength was set to 150 mM, and ΔG values were decomposed on a per-residue basis for each trajectory. Treatment of manganese(II) in mmpbsa.py required modification of the mdread2.F90 source file to assign the GB radius of the element.

### FTO WT and mutant expression vector construction

A codon optimized gblock of a human FTO(32-505) construct was obtained from IDT and cloned into a pET28a bacterial expression vector with N-terminal 6xHis tag. The FTO single point mutations were introduced into the 6xHis-FTO(32-505) construct using whole plasmid PCR site-directed mutagenesis.

### Protein expression and purification

FTO WT and mutant expression vectors were transformed into *E. coli* BL21(DE3) cells. 20 mL of an overnight culture was added to 2 L of LB media for growth at 37 °C with shaking at 200 rpm. When OD600 reached ∼0.6, cells were induced with 1 mM isopropyl β-D-1-thiogalactopyranoside for 18 hours at 18°C with shaking at 200 rpm. Centrifugation at 4100 x g was used to harvest cells and pellets were flash frozen and stored at - 70°C. Pellets were thawed and resuspended in lysis buffer (25 mM Tris-Base pH 8.0, 300 mM NaCl, 5 mM 2-mercaptoethanol), lysed by sonication, and centrifuged at 24000 x g for 45 min. Soluble His-tagged FTO was purified using HisPur Ni-NTA affinity resin (ThermoFisher Scientific) at 4°C and eluted in 25 mM Tris-HCl pH 7.5, 300 mM NaCl, 400 mM imidazole. Protein was buffer exchanged into 25 mM Tris-HCl pH 7.5, 25 mM NaCl and then purified using ion exchange chromatography with a 5 mL HiTrapQ column (Cytiva) and linear gradient into 25 mM Tris-HCl pH 7.5, 1 M NaCl. Further purification was done with size-exclusion chromatography using a Superdex 200pg 16/60 gel filtration column (Cytiva) in 10 mM HEPES pH 7.0, 50 mM KCl. Protein fractions were visualized using SDS-PAGE and purified FTO fractions were combined, concentrated, flash frozen, and stored in size-exclusion buffer with 10% glycerol at -70°C.

### FTO demethylation assays

*In vitro* FTO demethylation activity assays were performed by incubating 2 µM substrate (m^7^Gppp(m^6^A_m_)pG, m7Gppp(m^6^A_m_)pCpUpGpG, or CUGG(m^6^A)CUGG) with either 1 µM FTO or 0.1 µM FTO in demethylation reaction buffer (50 mM HEPES pH 7.0, 150 mM KCl, 75 µM (NH_4_)_2_Fe(SO_4_)_2_·6H2O, 300 µM 2-oxoglutarate, and 2 mM ascorbic acid). Reactions were initiated by addition of a 2X substrate mixture in reaction buffer to a 2X enzyme and cofactor mixture in reaction buffer and mixed with pipetting. Reaction time points were quenched by addition of EDTA to a final concentration of 1 mM.

### RNA nucleoside analysis and quantification

Quenched reaction samples with substrate m^7^Gppp(m^6^A_m_)pCpUpGpG were first decapped using NEB mRNA decapping enzyme (M0608S) for 1 hour. All quenched reaction samples were digested with NEB Nucleoside Digestion Mix (M0649S) for 1 hour (CUGG(m^6^A)CUGG) or 18 hours (m^7^Gppp(m^6^A_m_)pG or m^7^Gppp(m^6^A_m_)pCpUpGpG). Following RNA digestion into single nucleosides, for reaction samples with 1 µM FTO, protein was precipitated by addition of 20% TCA. Samples were then centrifuged and supernatant was collected for LC-MS analysis. Analysis of demethylation status was performed on an Agilent Bio-Inert 1260 Infinity II UHPLC system with Infinity Lab LC/MSD iQ. Separation of individual RNA nucleosides was conducted using an Agilent Zorbax SB-Aq Rapid Resolution HD (2.1 × 100 mm, 1.8 µm particle size) using mobile phase containing 0.1% formic acid (A) and 100% acetonitrile, 0.1% formic acid (B) and then detected by mass spectrometry in positive ionization mode. The gradient was as follows for detection of m^6^A at 0.480 mL/min (m/z = 282, 3.5 min) and A (m/z = 268, 1.6 min): 0-1.0 min, 100% A; 1.0-4.25 min, to 85% A/15% B; 4.25-5.0 min, to 25% A/75% B; 5.0-7.0 min, 25% A/75% B; 7.0-7.1 min, to 100% B; 7.1-11.75 min, 100% B. The gradient was as follows for detection of m^6^A_m_ at 0.450 mL/min (m/z = 296, 3.65 min) and A_m_ (m/z = 282, 3.5 min): 0-1.0 min, 100% A; 1.0-1.1 min, to 40% A/60% B; 1.1-5.0 min, to 28% A/72% B; 5.0-5.1 min, to 100% B; 5.1-8.0 min, 100% B.

### RNA substrate preparation

9-mer m^6^A RNA, CUGG(m^6^A)CUGG, was synthesized by IDT. Cap m^6^A_m_ RNA substrates (m^7^Gppp(m^6^A_m_)G and m7Gppp(m^6^A_m_)pCpUpGpG) were synthesized as previously described for similar RNA cap molecules.^86^

## Acknowledgments

This work was supported by the US National Institutes of Health, National Institute of General Medical Sciences award R35GM143000 to JSM, a T32GM133395 CBI fellowship to BS, and award P20GM104316, which funded key instrumentation used in this study. The content is solely the responsibility of the authors and does not necessarily represent the official views of the National Institutes of Health. This work was also supported by the BioStore resource, made possible by the US National Institutes of Health through the Delaware IDeA Network of Biomedical Research Excellence, awards P20GM103446 and S10OD028725. R.M.M. was supported by the National Science Foundation through an undergraduate research experience under award CBET-2232718 to J.A.H.-P. This work was further supported by grant 2019/33/B/ST4/01843 from the National Science Centre, Poland, to JJ. Computer time on Delta at the University of Illinois at Urbana-Champaign and Anvil at Purdue University was provided by allocation BIO240029 from the Advanced Cyberinfrastructure Coordination Ecosystem: Services & Support (ACCESS) program, funded by the National Science Foundation through awards #2138259, #2138286, #2138307, #2137603, and #2138296.

## References

(1) Yue, Y.; Liu, J.; He, C. RNA N6-Methyladenosine Methylation in Post-Transcriptional Gene Expression Regulation. Genes Dev. 2015, 29 (13), 1343–1355. 10.1101/gad.262766.115.

(2) Zaccara, S.; Ries, R. J.; Jaffrey, S. R. Reading, Writing and Erasing mRNA Methylation. Nat. Rev. Mol. Cell Biol. 2019, 20 (10), 608–624. 10.1038/s41580-019-0168-5.

(3) Dominissini, D.; Moshitch-Moshkovitz, S.; Schwartz, S.; Salmon-Divon, M.; Ungar, L.; Osenberg, S.; Cesarkas, K.; Jacob-Hirsch, J.; Amariglio, N.; Kupiec, M.; Sorek, R.; Rechavi, G. Topology of the Human and Mouse m6A RNA Methylomes Revealed by m6A-Seq. Nature 2012, 485 (7397), 201–206. 10.1038/nature11112.

(4) Meyer, K. D.; Saletore, Y.; Zumbo, P.; Elemento, O.; Mason, C. E.; Jaffrey, S. R. Comprehensive Analysis of mRNA Methylation Reveals Enrichment in 3′ UTRs and near Stop Codons. Cell 2012, 149 (7), 1635–1646. 10.1016/j.cell.2012.05.003.

(5) Xiao, W.; Adhikari, S.; Dahal, U.; Chen, Y.-S.; Hao, Y.-J.; Sun, B.-F.; Sun, H.-Y.; Li, A.; Ping, X.-L.; Lai, W.-Y.; Wang, X.; Ma, H.-L.; Huang, C.-M.; Yang, Y.; Huang, N.; Jiang, G.-B.; Wang, H.-L.; Zhou, Q.; Wang, X.-J.; Zhao, Y.-L.; Yang, Y.-G. Nuclear m6A Reader YTHDC1 Regulates mRNA Splicing. Mol. Cell 2016, 61 (4), 507–519. 10.1016/j.molcel.2016.01.012.

(6) Ping, X.-L.; Sun, B.-F.; Wang, L.; Xiao, W.; Yang, X.; Wang, W.-J.; Adhikari, S.; Shi, Y.; Lv, Y.; Chen, Y.-S.; Zhao, X.; Li, A.; Yang, Y.; Dahal, U.; Lou, X.-M.; Liu, X.; Huang, J.; Yuan, W.-P.; Zhu, X.-F.; Cheng, T.; Zhao, Y.-L.; Wang, X.; Danielsen, J. M. R.; Liu, F.; Yang, Y.-G. Mammalian WTAP Is a Regulatory Subunit of the RNA N6-Methyladenosine Methyltransferase. Cell Res. 2014, 24 (2), 177–189. 10.1038/cr.2014.3.

(7) Zhao, X.; Yang, Y.; Sun, B.-F.; Shi, Y.; Yang, X.; Xiao, W.; Hao, Y.-J.; Ping, X.-L.; Chen, Y.-S.; Wang, W.-J.; Jin, K.-X.; Wang, X.; Huang, C.-M.; Fu, Y.; Ge, X.-M.; Song, S.-H.; Jeong, H. S.; Yanagisawa, H.; Niu, Y.; Jia, G.-F.; Wu, W.; Tong, W.-M.; Okamoto, A.; He, C.; Rendtlew Danielsen, J. M.; Wang, X.-J.; Yang, Y.-G. FTO-Dependent Demethylation of N6-Methyladenosine Regulates mRNA Splicing and Is Required for Adipogenesis. Cell Res. 2014, 24 (12), 1403–1419. 10.1038/cr.2014.151.

(8) Roundtree, I. A.; Luo, G.-Z.; Zhang, Z.; Wang, X.; Zhou, T.; Cui, Y.; Sha, J.; Huang, X.; Guerrero, L.; Xie, P.; He, E.; Shen, B.; He, C. YTHDC1 Mediates Nuclear Export of N6-Methyladenosine Methylated mRNAs. eLife 2017, 6, e31311. 10.7554/eLife.31311.

(9) Lesbirel, S.; Viphakone, N.; Parker, M.; Parker, J.; Heath, C.; Sudbery, I.; Wilson, S. A. The m6A-Methylase Complex Recruits TREX and Regulates mRNA Export. Sci. Rep. 2018, 8 (1), 13827. 10.1038/s41598-018-32310-8.

(10) Heilman, K. L.; Leach, R. A.; Tuck, M. T. Internal 6-Methyladenine Residues Increase the in Vitro Translation Efficiency of Dihydrofolate Reductase Messenger RNA. Int. J. Biochem. Cell Biol. 1996, 28 (7), 823–829. 10.1016/1357-2725(96)00014-3.

(11) Wang, X.; Zhao, B. S.; Roundtree, I. A.; Lu, Z.; Han, D.; Ma, H.; Weng, X.; Chen, K.; Shi, H.; He, C. N6-Methyladenosine Modulates Messenger RNA Translation Efficiency. Cell 2015, 161 (6), 1388–1399. 10.1016/j.cell.2015.05.014.

(12) Meyer, K. D. m6A-Mediated Translation Regulation. Biochim. Biophys. Acta Gene Regul. Mech. 2019, 1862 (3), 301–309. 10.1016/j.bbagrm.2018.10.006.

(13) Shan, T.; Liu, F.; Wen, M.; Chen, Z.; Li, S.; Wang, Y.; Cheng, H.; Zhou, Y. m6A Modification Negatively Regulates Translation by Switching mRNA from Polysome to P-Body via IGF2BP3. Mol. Cell 2023, 83 (24), 4494–4508.e6. 10.1016/j.molcel.2023.10.040.

(14) Wang, X.; Lu, Z.; Gomez, A.; Hon, G. C.; Yue, Y.; Han, D.; Fu, Y.; Parisien, M.; Dai, Q.; Jia, G.; Ren, B.; Pan, T.; He, C. N6-Methyladenosine-Dependent Regulation of Messenger RNA Stability. Nature 2014, 505 (7481), 117–120. 10.1038/nature12730.

(15) Slobodin, B.; Bahat, A.; Sehrawat, U.; Becker-Herman, S.; Zuckerman, B.; Weiss, A. N.; Han, R.; Elkon, R.; Agami, R.; Ulitsky, I.; Shachar, I.; Dikstein, R. Transcription Dynamics Regulate Poly(A) Tails and Expression of the RNA Degradation Machinery to Balance mRNA Levels. Mol. Cell 2020, 78 (3), 434–444.e5. 10.1016/j.molcel.2020.03.022.

(16) Lee, Y.; Choe, J.; Park, O. H.; Kim, Y. K. Molecular Mechanisms Driving mRNA Degradation by m6A Modification. Trends Genet. 2020, 36 (3), 177–188. 10.1016/j.tig.2019.12.007.

(17) Batista, P. J.; Molinie, B.; Wang, J.; Qu, K.; Zhang, J.; Li, L.; Bouley, D. M.; Lujan, E.; Haddad, B.; Daneshvar, K.; Carter, A. C.; Flynn, R. A.; Zhou, C.; Lim, K.-S.; Dedon, P.; Wernig, M.; Mullen, A. C.; Xing, Y.; Giallourakis, C. C.; Chang, H. Y. m6A RNA Modification Controls Cell Fate Transition in Mammalian Embryonic Stem Cells. Cell Stem Cell 2014, 15 (6), 707–719. 10.1016/j.stem.2014.09.019.

(18) He, L.; Li, H.; Wu, A.; Peng, Y.; Shu, G.; Yin, G. Functions of N6-Methyladenosine and Its Role in Cancer. Mol. Cancer 2019, 18 (1), 176. 10.1186/s12943-019-1109-9.

(19) Fitzsimmons, C. M.; Batista, P. J. It’s Complicated… m6A-Dependent Regulation of Gene Expression in Cancer. Biochim. Biophys. Acta BBA - Gene Regul. Mech. 2019, 1862 (3), 382–393. 10.1016/j.bbagrm.2018.09.010.

(20) Deng, X.; Qing, Y.; Horne, D.; Huang, H.; Chen, J. The Roles and Implications of RNA m6A Modification in Cancer. Nat. Rev. Clin. Oncol. 2023, 20 (8), 507–526. 10.1038/s41571-023-00774-x.

(21) Wei, C.-M.; Gershowitz, A.; Moss, B. N6, O2′-Dimethyladenosine a Novel Methylated Ribonucleoside next to the 5′ Terminal of Animal Cell and Virus mRNAs. Nature 1975, 257 (5523), 251–253. 10.1038/257251a0.

(22) Akichika, S.; Hirano, S.; Shichino, Y.; Suzuki, T.; Nishimasu, H.; Ishitani, R.; Sugita, A.; Hirose, Y.; Iwasaki, S.; Nureki, O.; Suzuki, T. Cap-Specific Terminal N6-Methylation of RNA by an RNA Polymerase II–Associated Methyltransferase. Science 2019, 363 (6423). 10.1126/science.aav0080.

(23) Boulias, K.; Toczydłowska-Socha, D.; Hawley, B. R.; Liberman, N.; Takashima, K.; Zaccara, S.; Guez, T.; Vasseur, J.-J.; Debart, F.; Aravind, L.; Jaffrey, S. R.; Greer, E. L. Identification of the m6Am Methyltransferase PCIF1 Reveals the Location and Functions of m6Am in the Transcriptome. Mol. Cell 2019, 75 (3), 631–643.e8. 10.1016/j.molcel.2019.06.006.

(24) Sendinc, E.; Valle-Garcia, D.; Dhall, A.; Chen, H.; Henriques, T.; Navarrete-Perea, J.; Sheng, W.; Gygi, S. P.; Adelman, K.; Shi, Y. PCIF1 Catalyzes m6Am mRNA Methylation to Regulate Gene Expression. Mol. Cell 2019, 75 (3), 620–630.e9. 10.1016/j.molcel.2019.05.030.

(25) Sun, H.; Zhang, M.; Li, K.; Bai, D.; Yi, C. Cap-Specific, Terminal N6-Methylation by a Mammalian m6Am Methyltransferase. Cell Res. 2019, 29 (1), 80–82. 10.1038/s41422-018-0117-4.

(26) Mauer, J.; Luo, X.; Blanjoie, A.; Jiao, X.; Grozhik, A. V.; Patil, D. P.; Linder, B.; Pickering, B. F.; Vasseur, J.-J.; Chen, Q.; Gross, S. S.; Elemento, O.; Debart, F.; Kiledjian, M.; Jaffrey, S. R. Reversible Methylation of m6Am in the 5′ Cap Controls mRNA Stability. Nature 2017, 541 (7637), 371–375. 10.1038/nature21022.

(27) Mauer, J.; Sindelar, M.; Despic, V.; Guez, T.; Hawley, B. R.; Vasseur, J.-J.; Rentmeister, A.; Gross, S. S.; Pellizzoni, L.; Debart, F.; Goodarzi, H.; Jaffrey, S. R. FTO Controls Reversible m6Am RNA Methylation during snRNA Biogenesis. Nat. Chem. Biol. 2019, 15 (4), 340–347. 10.1038/s41589-019-0231-8.

(28) Chen, H.; Gu, L.; Orellana, E. A.; Wang, Y.; Guo, J.; Liu, Q.; Wang, L.; Shen, Z.; Wu, H.; Gregory, R. I.; Xing, Y.; Shi, Y. METTL4 Is an snRNA m6Am Methyltransferase That Regulates RNA Splicing. Cell Res. 2020, 30 (6), 544–547. 10.1038/s41422-019-0270-4.

(29) Sikorski, P. J.; Warminski, M.; Kubacka, D.; Ratajczak, T.; Nowis, D.; Kowalska, J.; Jemielity, J. The Identity and Methylation Status of the First Transcribed Nucleotide in Eukaryotic mRNA 5’ Cap Modulates Protein Expression in Living Cells. Nucleic Acids Res. 2020, 48 (4), 1607–1626. 10.1093/nar/gkaa032.

(30) Ben-Haim, M. S.; Pinto, Y.; Moshitch-Moshkovitz, S.; Hershkovitz, V.; Kol, N.; Diamant-Levi, T.; Beeri, M. S.; Amariglio, N.; Cohen, H. Y.; Rechavi, G. Dynamic Regulation of N6,2’-O-Dimethyladenosine (m6Am) in Obesity. Nat. Commun. 2021, 12 (1), 7185. 10.1038/s41467-021-27421-2.

(31) Zhuo, W.; Sun, M.; Wang, K.; Zhang, L.; Li, K.; Yi, D.; Li, M.; Sun, Q.; Ma, X.; Liu, W.; Teng, L.; Yi, C.; Zhou, T. m6Am Methyltransferase PCIF1 Is Essential for Aggressiveness of Gastric Cancer Cells by Inhibiting TM9SF1 mRNA Translation. Cell Discov. 2022, 8 (1), 48. 10.1038/s41421-022-00395-1.

(32) Jin, H.; Shi, Z.; Zhou, T.; Xie, S. Regulation of m6Am RNA Modification and Its Implications in Human Diseases. J. Mol. Cell Biol. 2024, 16 (3), mjae012. 10.1093/jmcb/mjae012.

(33) Liu, J. F.; Hawley, B. R.; Nicholson, L. S.; Jaffrey, S. R. Decoding m6Am by Simultaneous Transcription-Start Mapping and Methylation Quantification. eLife 2025, 13, RP104139. 10.7554/eLife.104139.

(34) Gerken, T.; Girard, C. A.; Tung, Y.-C. L.; Webby, C. J.; Saudek, V.; Hewitson, K. S.; Yeo, G. S. H.; McDonough, M. A.; Cunliffe, S.; McNeill, L. A.; Galvanovskis, J.; Rorsman, P.; Robins, P.; Prieur, X.; Coll, A. P.; Ma, M.; Jovanovic, Z.; Farooqi, I. S.; Sedgwick, B.; Barroso, I.; Lindahl, T.; Ponting, C. P.; Ashcroft, F. M.; O’Rahilly, S.; Schofield, C. J. The Obesity-Associated FTO Gene Encodes a 2-Oxoglutarate-Dependent Nucleic Acid Demethylase. Science 2007, 318 (5855), 1469–1472. 10.1126/science.1151710.

(35) Jia, G.; Fu, Y.; Zhao, X.; Dai, Q.; Zheng, G.; Yang, Y.; Yi, C.; Lindahl, T.; Pan, T.; Yang, Y.-G.; He, C. N6-Methyladenosine in Nuclear RNA Is a Major Substrate of the Obesity-Associated FTO. Nat. Chem. Biol. 2011, 7 (12), 885–887. 10.1038/nchembio.687.

(36) Wei, J.; Liu, F.; Lu, Z.; Fei, Q.; Ai, Y.; He, P. C.; Shi, H.; Cui, X.; Su, R.; Klungland, A.; Jia, G.; Chen, J.; He, C. Differential m6A, m6Am, and m1A Demethylation Mediated by FTO in the Cell Nucleus and Cytoplasm. Mol. Cell 2018, 71 (6), 973–985.e5. 10.1016/j.molcel.2018.08.011.

(37) Zhang, X.; Wei, L.-H.; Wang, Y.; Xiao, Y.; Liu, J.; Zhang, W.; Yan, N.; Amu, G.; Tang, X.; Zhang, L.; Jia, G. Structural Insights into FTO’s Catalytic Mechanism for the Demethylation of Multiple RNA Substrates. Proc. Natl. Acad. Sci. U. S. A. 2019, 116 (8), 2919–2924. 10.1073/pnas.1820574116.

(38) Ke, S.; Pandya-Jones, A.; Saito, Y.; Fak, J. J.; Vågbø, C. B.; Geula, S.; Hanna, J. H.; Black, D. L.; Darnell, J. E.; Darnell, R. B. m6A mRNA Modifications Are Deposited in Nascent Pre-mRNA and Are Not Required for Splicing but Do Specify Cytoplasmic Turnover. Genes Dev. 2017, 31 (10), 990–1006. 10.1101/gad.301036.117.

(39) McIntyre, A. B. R.; Gokhale, N. S.; Cerchietti, L.; Jaffrey, S. R.; Horner, S. M.; Mason, C. E. Limits in the Detection of m6A Changes Using MeRIP/m6A-Seq. Sci. Rep. 2020, 10 (1), 6590. 10.1038/s41598-020-63355-3.

(40) Merkestein, M.; Laber, S.; McMurray, F.; Andrew, D.; Sachse, G.; Sanderson, J.; Li, M.; Usher, S.; Sellayah, D.; Ashcroft, F. M.; Cox, R. D. FTO Influences Adipogenesis by Regulating Mitotic Clonal Expansion. Nat. Commun. 2015, 6, 6792. 10.1038/ncomms7792.

(41) Bartosovic, M.; Molares, H. C.; Gregorova, P.; Hrossova, D.; Kudla, G.; Vanacova, S. N6-Methyladenosine Demethylase FTO Targets Pre-mRNAs and Regulates Alternative Splicing and 3′-End Processing. Nucleic Acids Res. 2017, 45 (19), 11356–11370. 10.1093/nar/gkx778.

(42) Ronkainen, J.; Huusko, T. J.; Soininen, R.; Mondini, E.; Cinti, F.; Mäkelä, K. A.; Kovalainen, M.; Herzig, K.-H.; Järvelin, M.-R.; Sebert, S.; Savolainen, M. J.; Salonurmi, T. Fat Mass- and Obesity-Associated Gene Fto Affects the Dietary Response in Mouse White Adipose Tissue. Sci. Rep. 2015, 5 (1), 9233. 10.1038/srep09233.

(43) Chang, J. Y.; Park, J. H.; Park, S. E.; Shon, J.; Park, Y. J. The Fat Mass- and Obesity-Associated (FTO) Gene to Obesity: Lessons from Mouse Models. Obesity 2018, 26 (11), 1674–1686. 10.1002/oby.22301.

(44) Chen, A.; Chen, X.; Cheng, S.; Shu, L.; Yan, M.; Yao, L.; Wang, B.; Huang, S.; Zhou, L.; Yang, Z.; Liu, G. FTO Promotes SREBP1c Maturation and Enhances CIDEC Transcription during Lipid Accumulation in HepG2 Cells. Biochim. Biophys. Acta BBA - Mol. Cell Biol. Lipids 2018, 1863 (5), 538–548. 10.1016/j.bbalip.2018.02.003.

(45) Huang, C.; Chen, W.; Wang, X. Studies on the Fat Mass and Obesity-Associated (FTO) Gene and Its Impact on Obesity-Associated Diseases. Genes Dis. 2022, 10 (6), 2351–2365. 10.1016/j.gendis.2022.04.014.

(46) Poosri, S.; Boonyuen, U.; Chupeerach, C.; Soonthornworasiri, N.; Kwanbunjan, K.; Prangthip, P. Association of FTO Variants Rs9939609 and Rs1421085 with Elevated Sugar and Fat Consumption in Adult Obesity. Sci. Rep. 2024, 14 (1), 25618. 10.1038/s41598-024-77004-6.

(47) Boissel, S.; Reish, O.; Proulx, K.; Kawagoe-Takaki, H.; Sedgwick, B.; Yeo, G. S. H.; Meyre, D.; Golzio, C.; Molinari, F.; Kadhom, N.; Etchevers, H. C.; Saudek, V.; Farooqi, I. S.; Froguel, P.; Lindahl, T.; O’Rahilly, S.; Munnich, A.; Colleaux, L. Loss-of-Function Mutation in the Dioxygenase-Encoding FTO Gene Causes Severe Growth Retardation and Multiple Malformations. Am. J. Hum. Genet. 2009, 85 (1), 106–111. 10.1016/j.ajhg.2009.06.002.

(48) Li, L.; Zang, L.; Zhang, F.; Chen, J.; Shen, H.; Shu, L.; Liang, F.; Feng, C.; Chen, D.; Tao, H.; Xu, T.; Li, Z.; Kang, Y.; Wu, H.; Tang, L.; Zhang, P.; Jin, P.; Shu, Q.; Li, X. Fat Mass and Obesity-Associated (FTO) Protein Regulates Adult Neurogenesis. Hum. Mol. Genet. 2017, 26 (13), 2398–2411. 10.1093/hmg/ddx128.

(49) Wei, J.; Yu, X.; Yang, L.; Liu, X.; Gao, B.; Huang, B.; Dou, X.; Liu, J.; Zou, Z.; Cui, X.-L.; Zhang, L.-S.; Zhao, X.; Liu, Q.; He, P. C.; Sepich-Poore, C.; Zhong, N.; Liu, W.; Li, Y.; Kou, X.; Zhao, Y.; Wu, Y.; Cheng, X.; Chen, C.; An, Y.; Dong, X.; Wang, H.; Shu, Q.; Hao, Z.; Duan, T.; He, Y.-Y.; Li, X.; Gao, S.; Gao, Y.; He, C. FTO Mediates LINE1 m6A Demethylation and Chromatin Regulation in mESCs and Mouse Development. Science 2022, 376 (6596), 968–973. 10.1126/science.abe9582.

(50) Li, Z.; Weng, H.; Su, R.; Weng, X.; Zuo, Z.; Li, C.; Huang, H.; Nachtergaele, S.; Dong, L.; Hu, C.; Qin, X.; Tang, L.; Wang, Y.; Hong, G.-M.; Huang, H.; Wang, X.; Chen, P.; Gurbuxani, S.; Arnovitz, S.; Li, Y.; Li, S.; Strong, J.; Neilly, M. B.; Larson, R. A.; Jiang, X.; Zhang, P.; Jin, J.; He, C.; Chen, J. FTO Plays an Oncogenic Role in Acute Myeloid Leukemia as a N6-Methyladenosine RNA Demethylase. Cancer Cell 2017, 31 (1), 127–141. 10.1016/j.ccell.2016.11.017.

(51) Su, R.; Dong, L.; Li, Y.; Gao, M.; Han, L.; Wunderlich, M.; Deng, X.; Li, H.; Huang, Y.; Gao, L.; Li, C.; Zhao, Z.; Robinson, S.; Tan, B.; Qing, Y.; Qin, X.; Prince, E.; Xie, J.; Qin, H.; Li, W.; Shen, C.; Sun, J.; Kulkarni, P.; Weng, H.; Huang, H.; Chen, Z.; Zhang, B.; Wu, X.; Olsen, M. J.; Müschen, M.; Marcucci, G.; Salgia, R.; Li, L.; Fathi, A. T.; Li, Z.; Mulloy, J. C.; Wei, M.; Horne, D.; Chen, J. Targeting FTO Suppresses Cancer Stem Cell Maintenance and Immune Evasion. Cancer Cell 2020, 38 (1), 79–96.e11. 10.1016/j.ccell.2020.04.017.

(52) Relier, S.; Ripoll, J.; Guillorit, H.; Amalric, A.; Achour, C.; Boissière, F.; Vialaret, J.; Attina, A.; Debart, F.; Choquet, A.; Macari, F.; Marchand, V.; Motorin, Y.; Samalin, E.; Vasseur, J.-J.; Pannequin, J.; Aguilo, F.; Lopez-Crapez, E.; Hirtz, C.; Rivals, E.; Bastide, A.; David, A. FTO-Mediated Cytoplasmic m6Am Demethylation Adjusts Stem-like Properties in Colorectal Cancer Cell. Nat. Commun. 2021, 12 (1), 1716. 10.1038/s41467-021-21758-4.

(53) Zhang, K.; Zhang, F.; Wang, J. FTO Effects the Proliferation, Invasion, and Glycolytic Metabolism of Colon Cancer by Regulating PKM2. J. Cancer Res. Clin. Oncol. 2025, 151 (1), 36. 10.1007/s00432-024-06073-x.

(54) Xiao, L.; Li, X.; Mu, Z.; Zhou, J.; Zhou, P.; Xie, C.; Jiang, S. FTO Inhibition Enhances the Antitumor Effect of Temozolomide by Targeting MYC-miR-155/23a Cluster-MXI1 Feedback Circuit in Glioma. Cancer Res. 2020, 80 (18), 3945–3958. 10.1158/0008-5472.CAN-20-0132.

(55) Huff, S.; Tiwari, S. K.; Gonzalez, G. M.; Wang, Y.; Rana, T. M. m6A-RNA Demethylase FTO Inhibitors Impair Self-Renewal in Glioblastoma Stem Cells. ACS Chem. Biol. 2021, 16 (2), 324–333. 10.1021/acschembio.0c00841.

(56) Cui, Y.-H.; Yang, S.; Wei, J.; Shea, C. R.; Zhong, W.; Wang, F.; Shah, P.; Kibriya, M. G.; Cui, X.; Ahsan, H.; He, C.; He, Y.-Y. Autophagy of the m6A mRNA Demethylase FTO Is Impaired by Low-Level Arsenic Exposure to Promote Tumorigenesis. Nat. Commun. 2021, 12 (1), 2183. 10.1038/s41467-021-22469-6.

(57) Liu, Y.; Liang, G.; Xu, H.; Dong, W.; Dong, Z.; Qiu, Z.; Zhang, Z.; Li, F.; Huang, Y.; Li, Y.; Wu, J.; Yin, S.; Zhang, Y.; Guo, P.; Liu, J.; Xi, J. J.; Jiang, P.; Han, D.; Yang, C.-G.; Xu, M. M. Tumors Exploit FTO-Mediated Regulation of Glycolytic Metabolism to Evade Immune Surveillance. Cell Metab. 2021, 33 (6), 1221–1233.e11. 10.1016/j.cmet.2021.04.001.

(58) Duan, X.; Yang, L.; Wang, L.; Liu, Q.; Zhang, K.; Liu, S.; Liu, C.; Gao, Q.; Li, L.; Qin, G.; Zhang, Y. m6A Demethylase FTO Promotes Tumor Progression via Regulation of Lipid Metabolism in Esophageal Cancer. Cell Biosci. 2022, 12 (1), 60. 10.1186/s13578-022-00798-3.

(59) Garcia-Campos, M. A.; Edelheit, S.; Toth, U.; Safra, M.; Shachar, R.; Viukov, S.; Winkler, R.; Nir, R.; Lasman, L.; Brandis, A.; Hanna, J. H.; Rossmanith, W.; Schwartz, S. Deciphering the “m6A Code” via Antibody-Independent Quantitative Profiling. Cell 2019, 178 (3), 731–747.e16. 10.1016/j.cell.2019.06.013.

(60) Nicholson, L. S.; Guimarães-Teixeira, C.; Liu, J. F.; Poh, H. X.; Jaffrey, S. R. FTO Depletion Does Not Alter m6A Stoichiometry in AML mRNA: A Reassessment Using Direct RNA Nanopore Sequencing. bioRxiv: The Preprint Server for Biology October 23, 2025. 10.1101/2025.10.22.681652.

(61) Ashkenazy, H.; Abadi, S.; Martz, E.; Chay, O.; Mayrose, I.; Pupko, T.; Ben-Tal, N. ConSurf 2016: An Improved Methodology to Estimate and Visualize Evolutionary Conservation in Macromolecules. Nucleic Acids Res. 2016, 44 (W1), W344–W350. 10.1093/nar/gkw408.

(62) Yariv, B.; Yariv, E.; Kessel, A.; Masrati, G.; Chorin, A. B.; Martz, E.; Mayrose, I.; Pupko, T.; Ben-Tal, N. Using Evolutionary Data to Make Sense of Macromolecules with a “Face-Lifted” ConSurf. Protein Sci. 2023, 32 (3), e4582. 10.1002/pro.4582.

(63) Mazza, C.; Segref, A.; Mattaj, I. W.; Cusack, S. Large-scale Induced Fit Recognition of an m7GpppG Cap Analogue by the Human Nuclear Cap-binding Complex. EMBO J. 2002, 21 (20), 5548–5557. 10.1093/emboj/cdf538.

(64) Marcotrigiano, J.; Gingras, A.-C.; Sonenberg, N.; Burley, S. K. Cocrystal Structure of the Messenger RNA 5′ Cap-Binding Protein (eIF4E) Bound to 7-Methyl-GDP. Cell 1997, 89 (6), 951–961. 10.1016/S0092-8674(00)80280-9.

(65) Mugridge, J. S.; Tibble, R. W.; Ziemniak, M.; Jemielity, J.; Gross, J. D. Structure of the Activated Edc1-Dcp1-Dcp2-Edc3 mRNA Decapping Complex with Substrate Analog Poised for Catalysis. Nat. Commun. 2018, 9 (1), 1152. 10.1038/s41467-018-03536-x.

(66) Charenton, C.; Taverniti, V.; Gaudon-Plesse, C.; Back, R.; Séraphin, B.; Graille, M. Structure of the Active Form of Dcp1-Dcp2 Decapping Enzyme Bound to m7GDP and Its Edc3 Activator. Nat. Struct. Mol. Biol. 2016, 23 (11), 982–986. 10.1038/nsmb.3300.

(67) Eluwawalage, K. D. A.; Shimanski, B.; Warminski, M.; Katta, S.; Payne, R.; Yu, Y.; Kowalska, J.; Jemielity, J.; Mugridge, J. S. FTO Separation-of-Function Mutations Alter m6A versus m6Am Demethylation Selectivity on RNA. bioRxiv May 19, 2026, p 2026.05.19.726201. 10.64898/2026.05.19.726201.

(68) Jumper, J.; Evans, R.; Pritzel, A.; Green, T.; Figurnov, M.; Ronneberger, O.; Tunyasuvunakool, K.; Bates, R.; Žídek, A.; Potapenko, A.; Bridgland, A.; Meyer, C.; Kohl, S. A. A.; Ballard, A. J.; Cowie, A.; Romera-Paredes, B.; Nikolov, S.; Jain, R.; Adler, J.; Back, T.; Petersen, S.; Reiman, D.; Clancy, E.; Zielinski, M.; Steinegger, M.; Pacholska, M.; Berghammer, T.; Bodenstein, S.; Silver, D.; Vinyals, O.; Senior, A. W.; Kavukcuoglu, K.; Kohli, P.; Hassabis, D. Highly Accurate Protein Structure Prediction with AlphaFold. Nature 2021, 596 (7873), 583–589. 10.1038/s41586-021-03819-2.

(69) Case, D. A.; Aktulga, H. M.; Belfon, K.; Cerutti, D. S.; Cisneros, G. A.; Cruzeiro, V. W. D.; Forouzesh, N.; Giese, T. J.; Götz, A. W.; Gohlke, H.; Izadi, S.; Kasavajhala, K.; Kaymak, M. C.; King, E.; Kurtzman, T.; Lee, T.-S.; Li, P.; Liu, J.; Luchko, T.; Luo, R.; Manathunga, M.; Machado, M. R.; Nguyen, H. M.; O’Hearn, K. A.; Onufriev, A. V.; Pan, F.; Pantano, S.; Qi, R.; Rahnamoun, A.; Risheh, A.; Schott-Verdugo, S.; Shajan, A.; Swails, J.; Wang, J.; Wei, H.; Wu, X.; Wu, Y.; Zhang, S.; Zhao, S.; Zhu, Q.; Cheatham, T. E. I.; Roe, D. R.; Roitberg, A.; Simmerling, C.; York, D. M.; Nagan, M. C.; Merz, K. M. Jr. AmberTools. J. Chem. Inf. Model. 2023, 63 (20), 6183–6191. 10.1021/acs.jcim.3c01153.

(70) Zgarbová, M.; Otyepka, M.; Šponer, J.; Mládek, A.; Banáš, P.; Cheatham, T. E. I.; Jurečka, P. Refinement of the Cornell et al. Nucleic Acids Force Field Based on Reference Quantum Chemical Calculations of Glycosidic Torsion Profiles. J. Chem. Theory Comput. 2011, 7 (9), 2886–2902. 10.1021/ct200162x.

(71) Dolinsky, T. J.; Czodrowski, P.; Li, H.; Nielsen, J. E.; Jensen, J. H.; Klebe, G.; Baker, N. A. PDB2PQR: Expanding and Upgrading Automated Preparation of Biomolecular Structures for Molecular Simulations. Nucleic Acids Res. 2007, 35 (Web Server issue), W522–525. 10.1093/nar/gkm276.

(72) Humphrey, W.; Dalke, A.; Schulten, K. VMD: Visual Molecular Dynamics. J. Mol. Graph. 1996, 14 (1), 33–38. 10.1016/0263-7855(96)00018-5.

(73) Izadi, S.; Anandakrishnan, R.; Onufriev, A. V. Building Water Models: A Different Approach. J. Phys. Chem. Lett. 2014, 5 (21), 3863–3871. 10.1021/jz501780a.

(74) Tian, C.; Kasavajhala, K.; Belfon, K. A. A.; Raguette, L.; Huang, H.; Migues, A. N.; Bickel, J.; Wang, Y.; Pincay, J.; Wu, Q.; Simmerling, C. ff19SB: Amino-Acid-Specific Protein Backbone Parameters Trained against Quantum Mechanics Energy Surfaces in Solution. J. Chem. Theory Comput. 2020, 16 (1), 528–552. 10.1021/acs.jctc.9b00591.

(75) Li, P.; Merz, K. M. MCPB.Py: A Python Based Metal Center Parameter Builder. J. Chem. Inf. Model. 2016, 56 (4), 599–604. 10.1021/acs.jcim.5b00674.

(76) Case DA, Aktulga HM, Belfon K, Ben-Shalom IY, Berryman JT, Brozell SR, Carvalho FS, Cerutti DS, Cheatham TE III, Cisneros GA, Cruzeiro VWD, Darden TA, Forouzesh N, Ghazimirsaeed M, Giambasu G, Giese T, Gilson MK, Gohlke H, Goetz AW, Harris J, Huang Z, Izadi S, Izmailov SA, Kasavajhala K, Kaymak MC, Kolossváry I, Kovalenko A, Kurtzman T, Lee TS, Li P, Li Z, Lin C, Liu J, Luchko T, Luo R, Machado M, Manathunga M, Merz KM, Miao Y, Mikhailovskii O, Monard G, Nguyen H, O’Hearn KA, Onufriev A, Pan F, Pantano S, Rahnamoun A, Roe DR, Roitberg A, Sagui C, Schott-Verdugo S, Shajan A, Shen J, Simmerling CL, Skrynnikov NR, Smith J, Swails J, Walker RC, Wang J, Wang J, Wu X, Wu Y, Xiong Y, Xue Y, York DM, Zhao C, Zhu Q, Kollman PA. Amber 2025. University of California, San Francisco, 2025.

(77) Wang, J.; Wolf, R. M.; Caldwell, J. W.; Kollman, P. A.; Case, D. A. Development and Testing of a General Amber Force Field. J. Comput. Chem. 2004, 25 (9), 1157–1174. 10.1002/jcc.20035.

(78) Steinbrecher, T.; Latzer, J.; Case, D. A. Revised AMBER Parameters for Bioorganic Phosphates. J. Chem. Theory Comput. 2012, 8 (11), 4405–4412. 10.1021/ct300613v.

(79) Aduri, R.; Psciuk, B. T.; Saro, P.; Taniga, H.; Schlegel, H. B.; SantaLucia, J. AMBER Force Field Parameters for the Naturally Occurring Modified Nucleosides in RNA. J. Chem. Theory Comput. 2007, 3 (4), 1464–1475. 10.1021/ct600329w.

(80) D.A. Case, H.M. Aktulga, K. Belfon, I.Y. Ben-Shalom, J.T. Berryman, S.R. Brozell, F.S. Carvalho, D.S. Cerutti, T.E. Cheatham, III, G.A. Cisneros, V.W.D. Cruzeiro, T.A. Darden, N. Forouzesh, M. Ghazimirsaeed, G. Giambaşu, T. Giese, M.K. Gilson, H. Gohlke, A.W. Goetz, J. Harris, Z. Huang, S. Izadi, S.A. Izmailov, K. Kasavajhala, M.C. Kaymak, I. Kolossvary, A. Kovalenko, T. Kurtzman, T.S. Lee, P. Li, Z. Li, C. Lin, J. Liu, T. Luchko, R. Luo, M. Machado, M. Manathunga, K.M. Merz, Y. Miao, O. Mikhailovskii, G. Monard, H. Nguyen, K.A. O’Hearn, A. Onufriev, F. Pan, S. Pantano, A. Rahnamoun, D.R. Roe, A. Roitberg, C. Sagui, S. Schott-Verdugo, A. Shajan, J. Shen, C.L. Simmerling, N.R. Skrynnikov, J. Smith, J. Swails, R.C. Walker, J. Wang, J. Wang, X. Wu, Y. Wu, Y. Xiong, Y. Xue, D.M. York, C. Zhao, Q. Zhu, and P.A. Kollman. Amber 2024 2024, University of California, San Francisco.

(81) Sindhikara, D. J.; Kim, S.; Voter, A. F.; Roitberg, A. E. Bad Seeds Sprout Perilous Dynamics: Stochastic Thermostat Induced Trajectory Synchronization in Biomolecules. J. Chem. Theory Comput. 2009, 5 (6), 1624–1631. 10.1021/ct800573m.

(82) Srinivasan, J.; Cheatham, T. E.; Cieplak, P.; Kollman, P. A.; Case, D. A. Continuum Solvent Studies of the Stability of DNA, RNA, and Phosphoramidate–DNA Helices. J. Am. Chem. Soc. 1998, 120 (37), 9401–9409. 10.1021/ja981844+.

(83) Massova, I.; Kollman, P. A. Combined Molecular Mechanical and Continuum Solvent Approach (MM-PBSA/GBSA) to Predict Ligand Binding. Perspect. Drug Discov. Des. 2000, 18 (1), 113–135. 10.1023/A:1008763014207.

(84) Miller, B. R. I.; McGee, T. D. Jr.; Swails, J. M.; Homeyer, N.; Gohlke, H.; Roitberg, A. E. MMPBSA.Py: An Efficient Program for End-State Free Energy Calculations. J. Chem. Theory Comput. 2012, 8 (9), 3314–3321. 10.1021/ct300418h.

(85) Onufriev, A.; Bashford, D.; Case, D. A. Exploring Protein Native States and Large-Scale Conformational Changes with a Modified Generalized Born Model. Proteins 2004, 55 (2), 383–394. 10.1002/prot.20033.

(86) Warminski, M.; Trepkowska, E.; Smietanski, M.; Sikorski, P. J.; Baranowski, M. R.; Bednarczyk, M.; Kedzierska, H.; Majewski, B.; Mamot, A.; Papiernik, D.; Popielec, A.; Serwa, R. A.; Shimanski, B. A.; Sklepkiewicz, P.; Sklucka, M.; Sokolowska, O.; Spiewla, T.; Toczydlowska-Socha, D.; Warminska, Z.; Wolosewicz, K.; Zuberek, J.; Mugridge, J. S.; Nowis, D.; Golab, J.; Jemielity, J.; Kowalska, J. Trinucleotide mRNA Cap Analogue N6-Benzylated at the Site of Posttranscriptional m6Am Mark Facilitates mRNA Purification and Confers Superior Translational Properties In Vitro and In Vivo. J. Am. Chem. Soc. 2024, 146 (12), 8149–8163. 10.1021/jacs.3c12629.

