## Supplementary Information for "RNA demethylase FTO uses conserved aromatic residues to recognize the mRNA 5′ cap and promote efficient m^6^A_m_ demethylation"

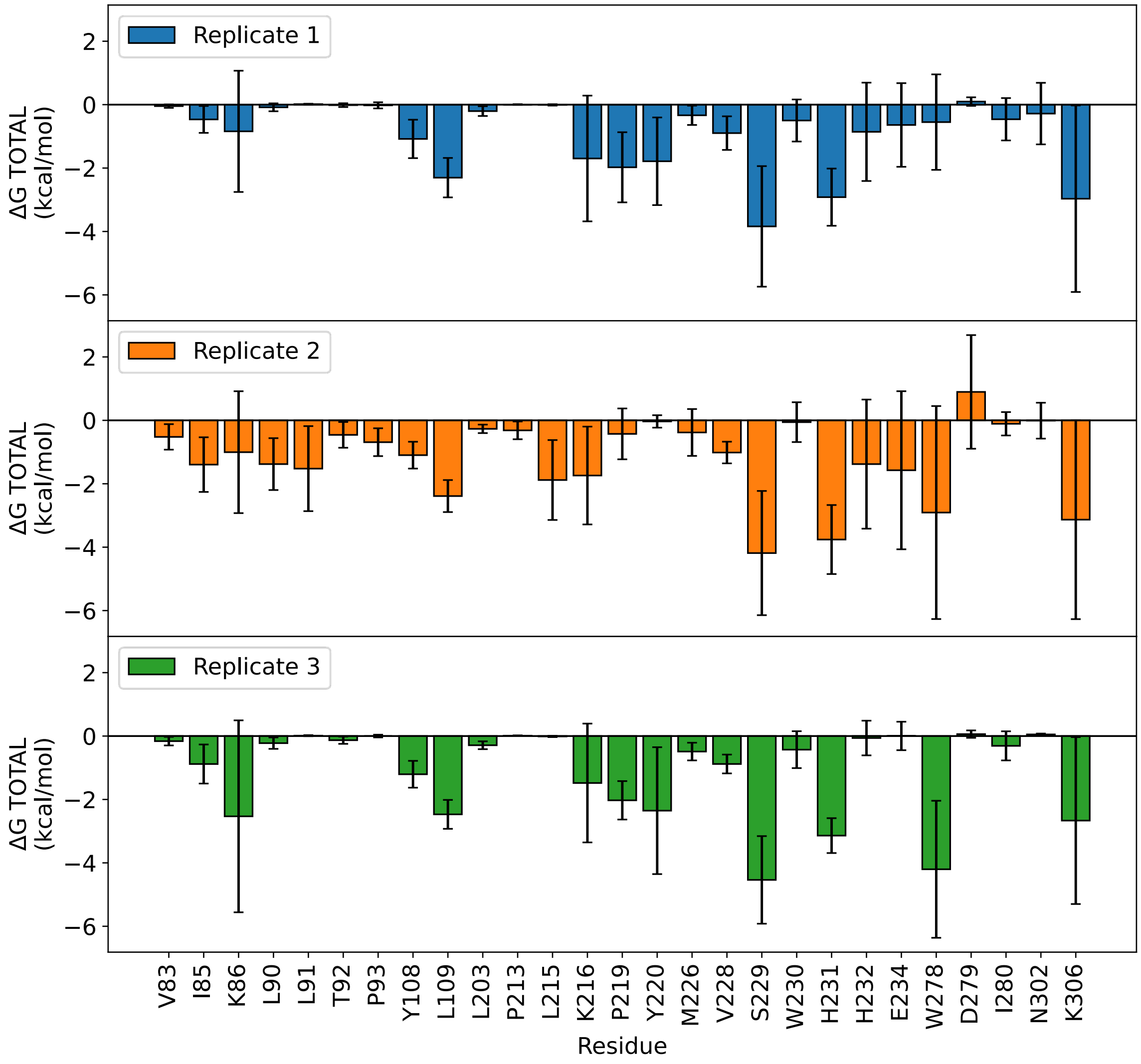


**Supplementary Figure 1. FTO-$m^7$Gppp($m^6A_m$)pG binding energies determined from MD simulations.** Average per-residue contributions to the total Gibbs free energy of binding **(**ΔG, kcal/mol) calculated using the Molecular Mechanics/Generalized Born Surface Area (MM/GBSA) method. Error bars represent standard deviation. FTO residues are included here if the magnitude of their contribution was >0.25 kcal/mol in at least one replicate. Negative values indicate favorable contributions to binding. Values are averaged over one million MD frames per replicate.

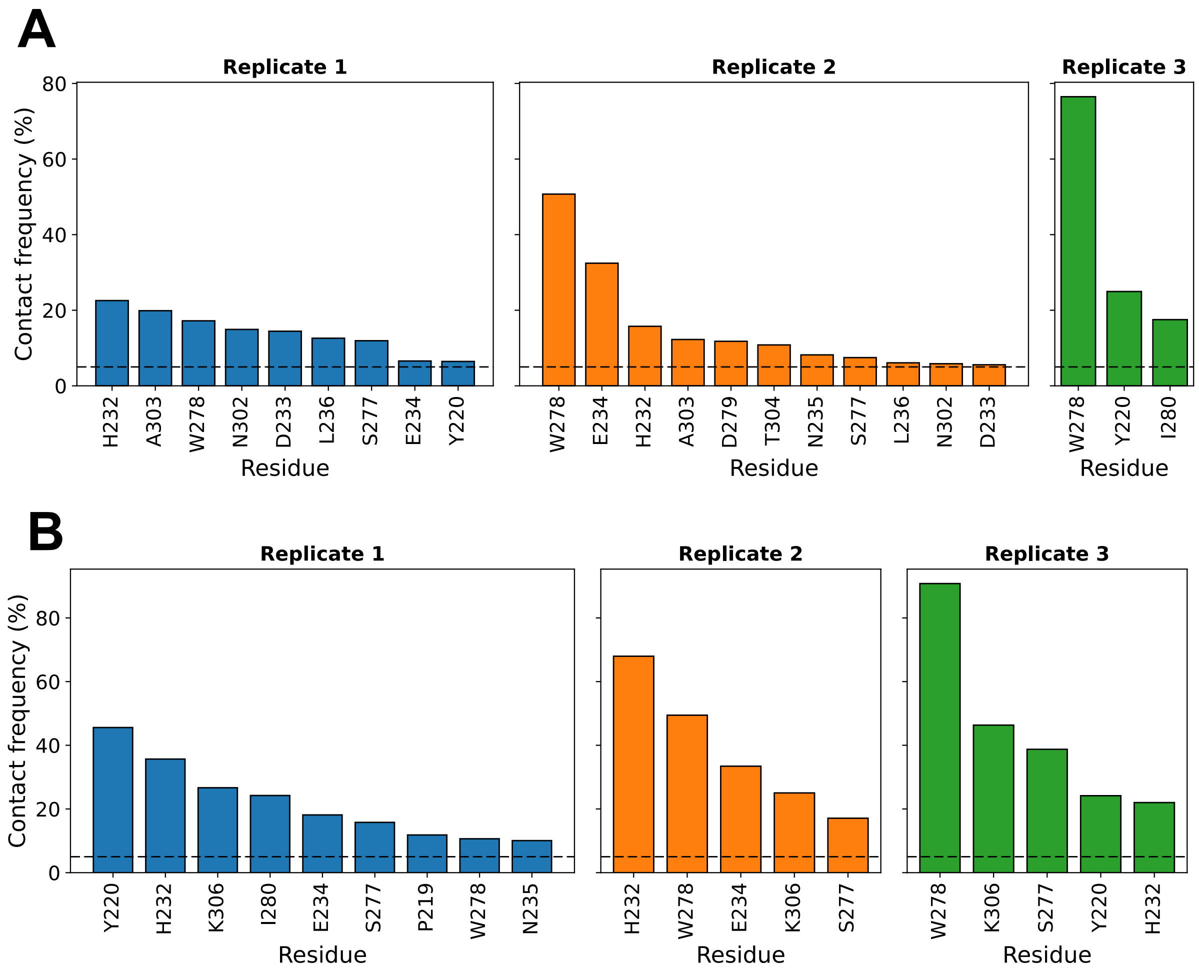


**Supplementary Figure 2. FTO-m$^7$G contact frequencies determined from MD simulations.** Contact between FTO residues and m$^7$G was defined as atomic distances ≤ 3.5 Å. FTO residues are included here if their m$^7$G contact frequency exceeded 5% of MD trajectory frames; percentages were calculated out of one million frames per replicate. Panel A lists FTO residues contacting the m$^7$G base, and panel B lists FTO residues contacting the m$^7$G ribose for each MD replicate.

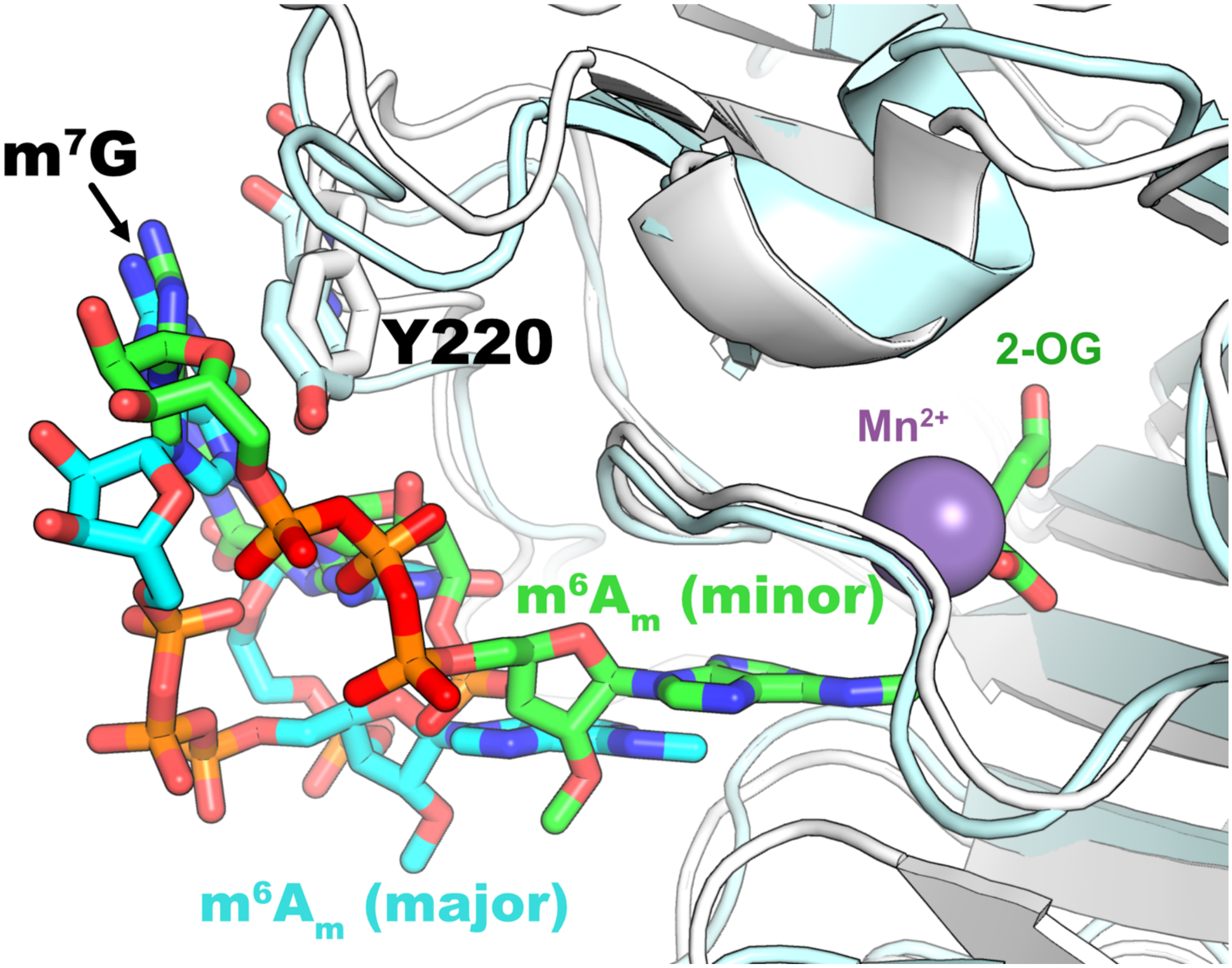


**Supplementary Figure 3. MD simulations suggest that Y220-m$^7$G base stacking is not conducive to m$^6$A$_m$ demethylation.** During a preliminary run of Replicate 2, m$^7$G lost contact with FTO residues, leading to destabilization of the m7Gppp(m6Am)pG binding mode and shifting of the m$^6$A$_m$ base away from the active site metal. The MD simulation was restarted from a frame in which appropriate m$^7$G and m$^6$A$_m$ interactions remained in place to continue sampling the catalytically relevant complex. However, the preliminary run demonstrated that Y220-m$^7$G base stacking was favored when the m$^6$A$_m$ base was only shallowly positioned in the active site (cyan 'major' conformation above, in which the m$^6$A methyl group is ~8 Å from active site metal; corresponds to light cyan FTO cartoon). A limited number of trajectory frames preceding the ligand shift showed m$^7$G-Y220 interaction modes with the m$^6$A$_m$ base positioned deeper within the active site, in a conformation more consistent with demethylation catalysis (green 'minor' conformation above, in which the m$^6$A methyl group is ~4 Å from active site metal; corresponding to white FTO cartoon). These results suggest that Y220-m$^7$G base stacking may promote m$^6$A$_m$ cap binding modes that are ineffective for catalysis.

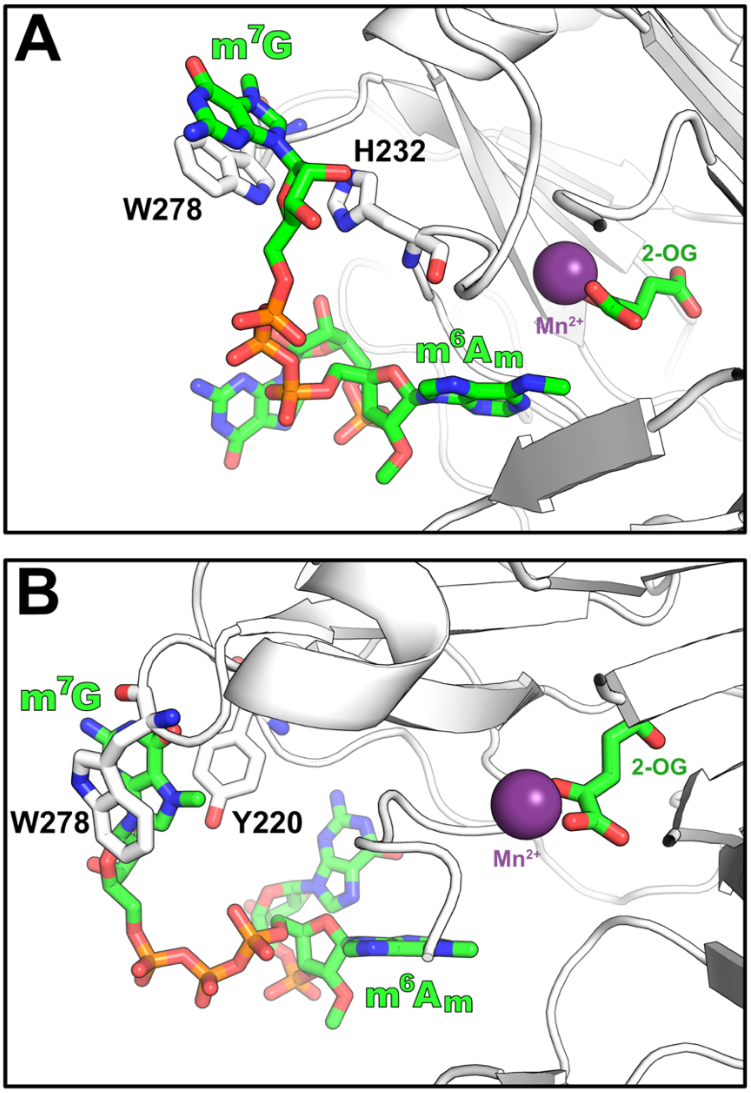


**Supplementary Figure 4. FTO-m$^7$G binding modes with dual interactions observed in MD simulations**. **(A)** Dual interaction mode where H232 engages the m$^7$G ribose through CH-π/van der Waals interactions, and W278 stacks or forms cation-π interactions with the m$^7$G guanosine base. **(B)** Dual interaction mode observed where Y220 and W278 engage the m$^7$G base in simultaneous stacking / cation-π interactions, which correlates with the m$^6$A$_m$ base being withdrawn somewhat from the active site pocket. In this particular MD frame, the m$^6$A methyl group is ~10 Å from the active site metal. The latter mode was observed during a preliminary run of Replicate 2, in which m$^7$G lost contact with FTO residues, leading to destabilization of the m$^7$Gppp(m$^6$A$_m$)pG binding mode and shifting of the m$^6$A$_m$ base away from the active site metal. The MD simulation was restarted from a frame in which appropriate m$^7$G and m$^6$A$_m$ interactions remained in place to continue sampling the catalytically relevant complex. However, the preliminary run demonstrated that Y220-m$^7$G base stacking was favored only with m$^6$A$_m$ cap binding modes expected to be ineffective for catalysis.

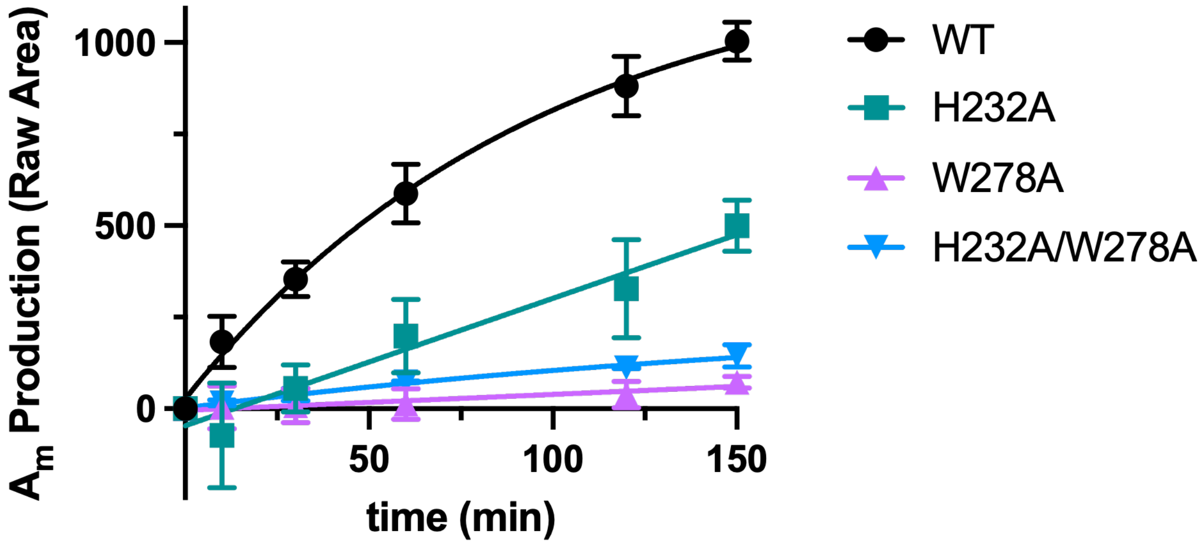


**Supplementary Figure 5. $m^6A_m$ demethylation time courses with cap substrates.** Mutation of conserved aromatic residues in FTO identified as $m^7G$ interaction sites show reduced production of $A_m$ product during demethylation reactions with model $m^6A_m$ cap substrates. Demethylation time courses were conducted with 2 μM $m^7Gppp(m^6A_m)pG$ and 1 μM FTO enzyme in triplicate. At various time points from 0 to 150 minutes, reaction samples were quenched with EDTA, digested to single nucleosides with RNAase cocktail, and the relative amounts of $A_m$ were quantified by UHPLC-MS. The integrated MS area (raw area) is plotted versus time and data are shown as mean values ± SEM (n = 3).

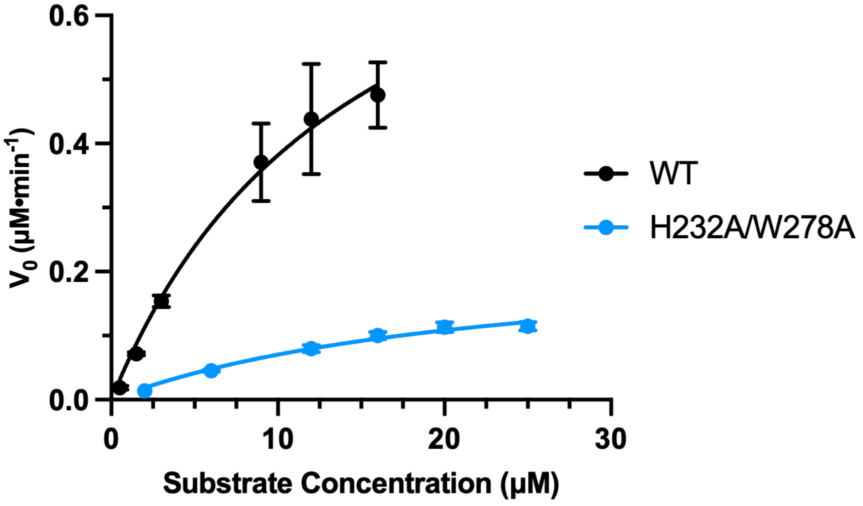


**Supplementary Figure 6. Michaelis-Menten analysis of $m^6A_m$ demethylation by FTO WT vs H232A/W278A mutant**. Michaelis-Menten kinetic plots for FTO-mediated demethylation of $m^7$Gppp($m^6A_m$)pCpUpGpG cap substrate by WT versus H232A/W278A double mutant. For WT FTO, $k_{cat}$ = 4.7 ± 1.6 $min^{-1}$ and $K_M$ = 15 ± 9 µM; for H232A/W278A FTO, $k_{cat}$ = 1.2 ± 0.2 $min^{-1}$ and $K_M$ = 23 ± 7 µM. This corresponds to a ~4-fold change in $k_{cat}$, and a less than 2-fold change in $K_M$ where the determined $K_M$ values are within error of one another, suggesting these FTO mutations predominantly impact $k_{cat}$. WT FTO kinetic data are as previously reported in Calzini et al, *bioRxiv* **2026**, *doi*:10.1101/2025.05.06.652568.

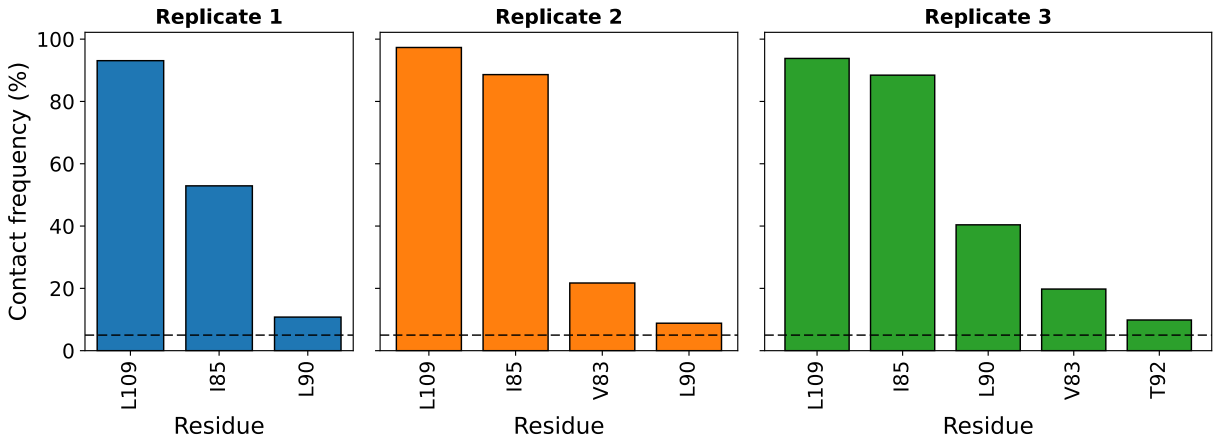


**Supplementary Figure 7. FTO-$A_m$ 2′-O-methyl group contact frequencies determined from MD simulations.** Contact between FTO residues and the methyl group was defined as atomic distances ≤ 3.5 Å. FTO residues are included here if their methyl group contact frequency exceeded 5% of MD trajectory frames; percentages were calculated out of one million frames per replicate.

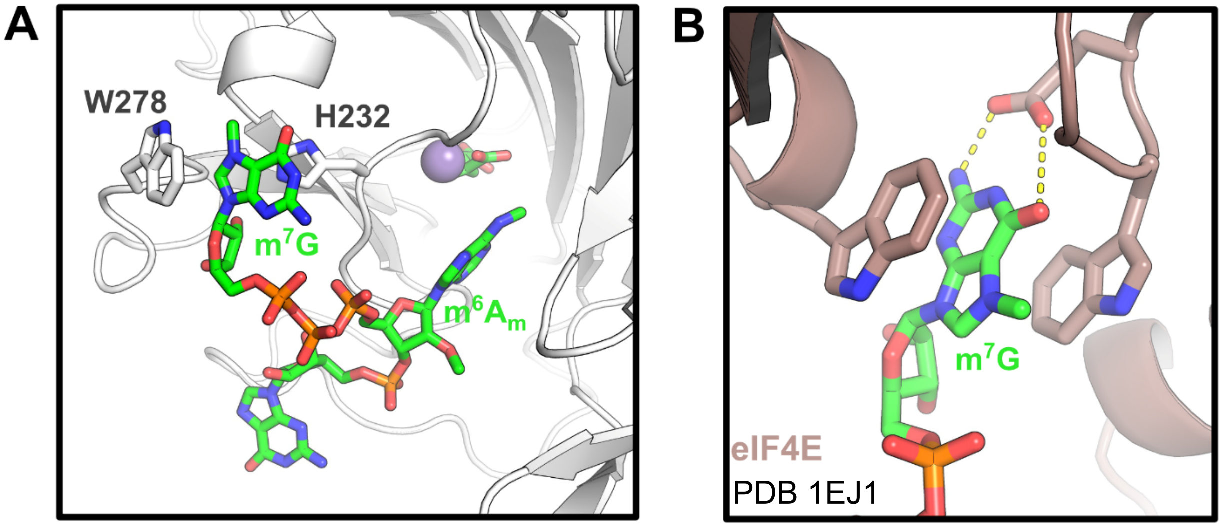


**Supplementary Figure 8. Simultaneous stacking interaction of aromatic residues with $m^7G$. (A)** Representative MD frame showing $m^7G$ weakly stacked between conserved aromatic FTO residues H232 and W278, which was observed only rarely during MD simulations. **(B)** Canonical cap recognition showing $m^7G$ stacked between two aromatic residues of eIF4E from PDB 1EJ1.
